# Notch engagement by Jag1 nanoscale clusters indicates a force-independent mode of activation

**DOI:** 10.1101/2022.11.22.517517

**Authors:** Ioanna Smyrlaki, Ferenc Fördös, Iris Rocamonde Lago, Yang Wang, Antonio Lentini, Vincent C. Luca, Björn Reinius, Ana I. Teixeira, Björn Högberg

## Abstract

The Notch signaling pathway is a cell-cell communication system with fundamental roles in embryonic development and the nervous system. The model of Notch receptor activation that is currently most accepted, involves a force-induced conformation change at the negative regulatory region of the receptor, the subsequent recruitment of ADAM metalloproteases and a cleavage cascade that releases the Notch intracellular domain. Here, we define conditions that enable force-independent Notch activation through the formation of soluble, long-lived, multivalent ligand-receptor complexes. To investigate how ligand valency affects activation of Notch receptors, we treated iPSc-derived neuroepithelial stem-like (lt-NES) cells with different spatially defined, molecularly precise ligand nanopatterns on DNA origami nanostructures. Our data indicate that Notch signaling is activated via stimulation with multivalent clusters of the ligand Jag1, and even multivalent chimeric structures where some Jag1 proteins are replaced by other binders that do not target Notch. The findings are corroborated by systematic elimination, through experimental control, of several confounding factors that potentially could generate forces, including electrostatic interactions, endocytosis and non-specific binding. Taken together, our data suggest a model where Jag1 ligands are able to activate Notch receptors upon prolonged binding, which subsequently triggers downstream signaling in a force independent manner. These findings reveal a distinct mode of activation of Notch and could lay the foundation for the development of soluble Notch agonists that currently remain elusive.

## INTRODUCTION

Notch signaling is an evolutionary conserved cell-to-cell communication system present in most animals with fundamental roles in cell fate decisions, tissue patterning, angiogenesis, and neurogenesis. In mammals, four types of Notch receptors (Notch1-4) are present and two families of ligands, Serrate (Jag1 and Jag2) and Delta (DLL1, DLL3 and DLL4)^1^. Activation of this pathway relies on three proteolytic steps performed on three cleavage sites in the Notch receptor (S1-S3) (Fig. 1B). During its maturation in the Golgi apparatus, the Notch receptor is cleaved by furin-like convertase (S1) forming a non-covalently associated heterodimer.^2,3^ The interaction between a receptor at the cell surface and a ligand expressed on an adjacent cell, leads to an extracellular ADAM mediated cleavage (S2) and a transmembrane γ-secretase (S3) mediated cut, resulting in release of the Notch intracellular domain (NTCD).^4,5^ After these cleavages, the NTCD translocates to the nucleus to form a complex with a DNA binding protein of CSL family and the co-activator Mastermind, together acting as a transcriptional activator complex^6^.

**Figure 1:**
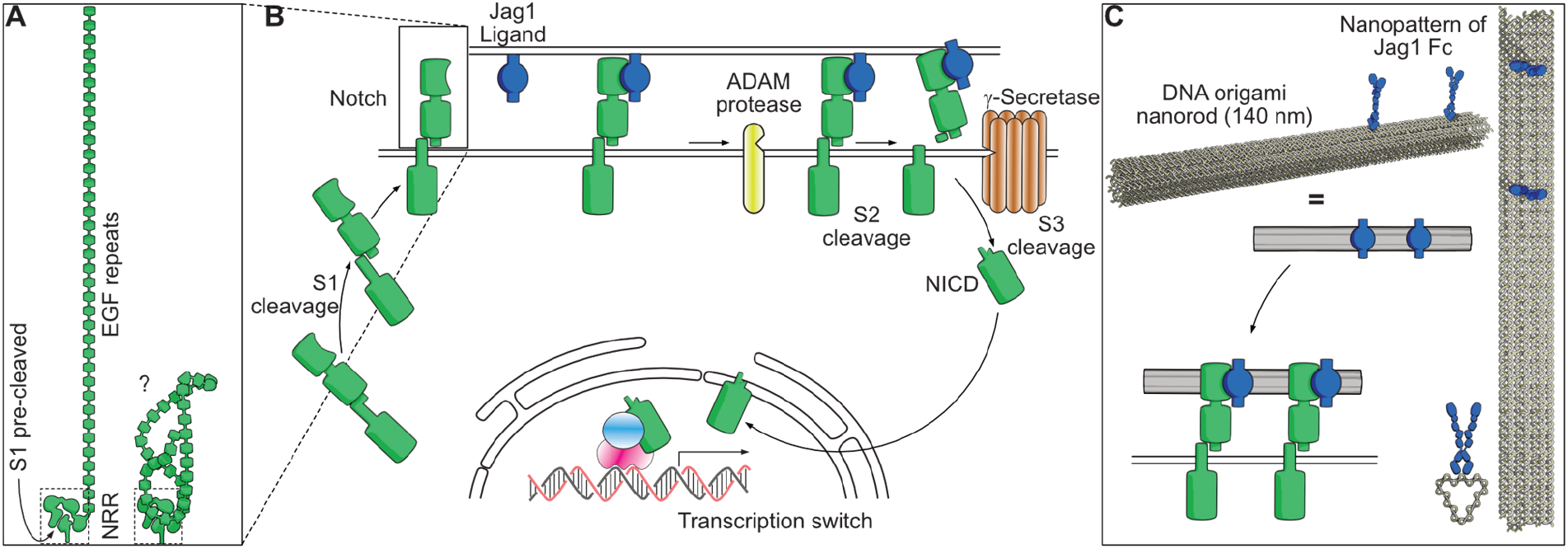
Schematic view of the Notch pathway and experimental principle. (**A**) **The extracellular domain of the Notch receptor** comprises a negatively regulatory region (NRR, dashed box) containing Lin12-Notch repeats and a pre-cleaved S1 site held together non-covalently. The most prominent feature of the receptor is its 29-36 (depending on Notch subtype) EGF repeats. Whether these are stretched out (left) or forms some tertiary structure (right), is not fully understood. **(B) The canonical Notch pathway** from maturation where Notch heterodimers are formed via Furin cleavage of the S1 in the Golgi, followed by membrane insertion. Binding to ligands (in this case Jag1) induces cleavage of S2 by ADAM proteases, which in turn enables cleavage of S3 via γ-Secretase to release the Notch intracellular domain (NTCD). The NTCD translocates to the nucleus where it forms a transcription activator complex with CSL and Mastermind. **(C) DNA origami nanopatterns of Jag1** were designed where Jag1-Fc molecules covalently conjugated to DNA oligonucleotides could be attached to the nanostructure to form many types of precisely controlled multivalent Jag1 binder patterns.

In mammals, the extracellular part of the Notch receptors (NECD) is composed of 29-36 EGF repeats (Fig. 1A). The exact function of all of the repeats are not well understood but some appear necessary for the interactions with their ligands^7,8^ and others are thought to bind to calcium ions to potentially modulate the affinity of ligand binding.^9,10^ Whether the repeat region is rod-like (Fig. 1A, left) or if it could possess a folded tertiary structure (Fig. 1A, right) is unclear. Following the EGF repeats, are three cysteine-rich Lin12-Notch repeats (LNR) and a hydrophobic heterodimerization domain at the pre-cleaved S1 that together form what is known as the negative regulatory region (NRR)^11^. The NRR is located in the receptor close to the cell membrane and is hypothesized to form a loop that protects the S2 site from uncontrolled ADAM cleavage activity.^12–14^ Within the transmembrane segment, the S3 site serves as the key point for the NTCD release.^5,15^ The NTCD contains a RAM domain and seven ankyrin repeats that are involved in CLS interaction at the nucleus, a transcription activation domain (TAD) and a PEST region responsible for NTCD degradation.^16–20^

Ligands of Notch are also single pass transmembrane proteins^1^. The structure of Notch ligands of both families contains a DSL domain at the N-terminus and EGF-like repeats (16 for Jagged family and 5-9 for Delta family) followed by a cysteine rich domain (CRD) which is present only in Jagged family ligands. Further studies show that the N-terminus of human Jagged-1 and DLL-1 is a C2 phospholipid recognition domain that binds phospholipid bilayers in a Ca^2+^ dependent fashion.^10^ Co-crystal structures of Notch1 and DLL4 have revealed that the DSL and C2 domains of DLL4 bind to EGF11 and EGF12 of Notch1^8^ while the C2 and EGF3 of Jag1 engage EGF8 through EGF12, respectively.^21^ These data imply that different sites on the Notch receptors preferentially interact with specific ligands.

The current model of canonical Notch pathway activation requires Notch endocytosis by the ligand expressing cell.^22,23^ A more recent study suggests that the Epsin pathway is a crucial mediator for endocytosis of the complex, while engulfment of the ligand by receptor expressing cells, terminates the signal without leading to activation.^24^ The two dominant theories for this endocytosis are either (i) a receptor recycling model where each receptor (and ligand) should be used only once, or (ii) a pulling model in which a force would be required to uncover the S2 region of the receptor. Tension gauge tethers have been applied on ligands immobilized on surface, to show that 12pN force is sufficient for Notch activation.^25^ The model of activation by pulling force was further confirmed by magnetic beads tethered to Notch molecules on the cell surface.^26^ Another systematic study revealed that DLL4, which binds to Notch with higher affinity than Jag1, activates Notch at tensions even as low as 1pN while Jag1 would require tension above 4pN.^21^ In bio-membrane force probe experiments, where force clamp spectroscopy was used to measure lifetimes of receptor ligands bonds under a range of tensile forces, a catch bond behavior appears to occur for both Jag1 and DLL4.^27^ It is also clear, from the structure of the receptor, that excessive pulling^28^ or disruption of the heterodimerization domain by chelators^29^ will shed the ECD and expose the receptor for activation. Thus, while it appears clear that a pulling force can activate Notch, we cannot rule out that other modes of activation could be equally important if they were to occur in nature.

The effect of patterns of Notch ligands has previously been investigated at the microscale and on surfaces.^30,31^ However, the effect of nanoscale patterning of ligands, and in particular patterns displayed from a solution phase, has not been systematically evaluated. In this study we explore the hypothesis that clusters of ligands prolong the formation of ligand-receptor complexes, which in turn exposes the NRR region for ADAM10 mediated cleavage and subsequent activation of the Notch pathway. DNA origami^32,33^ has been demonstrated as a potent tool to finely control the nanoscale formation of ligand clusters with response on spatial distribution of receptors.^34,35,36^ Herein, we applied this technique to generate molecularly precise Jag1 nanopatterns (JNP) capable of stimulating Notch-expressing cells in solution (Fig 1C). We used long-term self-renewing neuroepithelial-like stem cells (lt-NES cells)^37^ as the cellular model for studying the effect of ligand multivalency. This system was chosen to give the best model available for endogenous Notch activity in cells that resemble the neuronal progenitors in the early neural tube.

We consistently found that larger Jag1 clusters induce higher levels of Notch activation. Moreover, this interaction appear to occur independently of pulling forces. Instead, our results are consistent with a model where the lifespan of ligand-receptor complex increases with multivalency and regulates Notch activation. Based on our findings, we hypothesize that prolonged receptor-ligand interactions trigger force-independent conformational changes in the receptor that expose the NRR to ADAM10 cleavage, leading to subsequent initiation of activation.

## RESULTS

### Achieving controlled Jag1 nanopatterns

We used a rod like DNA origami structure^34^ (Fig. 1C, Methods) to spatially position zero (baseline-control), one, two, three, four or eight, dimeric Jag1Fc fusion proteins (Fig. 2A) and study the ability of ligand clusters to activate the Notch signaling pathway. To assemble the complex of proteins with the DNA origami, we hybridized the folded DNA nanostructures, containing single strand protruding oligos at desired positions, to a complementary single strand DNA (ssDNA) oligo already conjugated to the Jag1Fc protein.^34^ To achieve site-specific conjugation of the protein to a ssDNA-anchor, we use a Bis-sulfon-DBCO bispecific crosslinking compound.^36^ First, we modified the protein with the compound by selective reaction between the bis-sulfon group and the histidines at the C terminus of the protein, and then an azide modified oligonucleotide was conjugated to the protein by click chemistry between the azide-and the DBCO-group. Using a gel retardation assay, we confirmed successful production of Jag1 nanopatterns (JNP), where an increasing shift was observed as more proteins were hybridized onto the DNA origami nanorod (Figure 2B). During the hybridization reaction of Jag1-ssDNA with the complementary overhangs on DNA nanostructures, we used an excess of Jag1 conjugates, which were later removed by size exclusion chromatography (see Methods).

**Figure 2:**
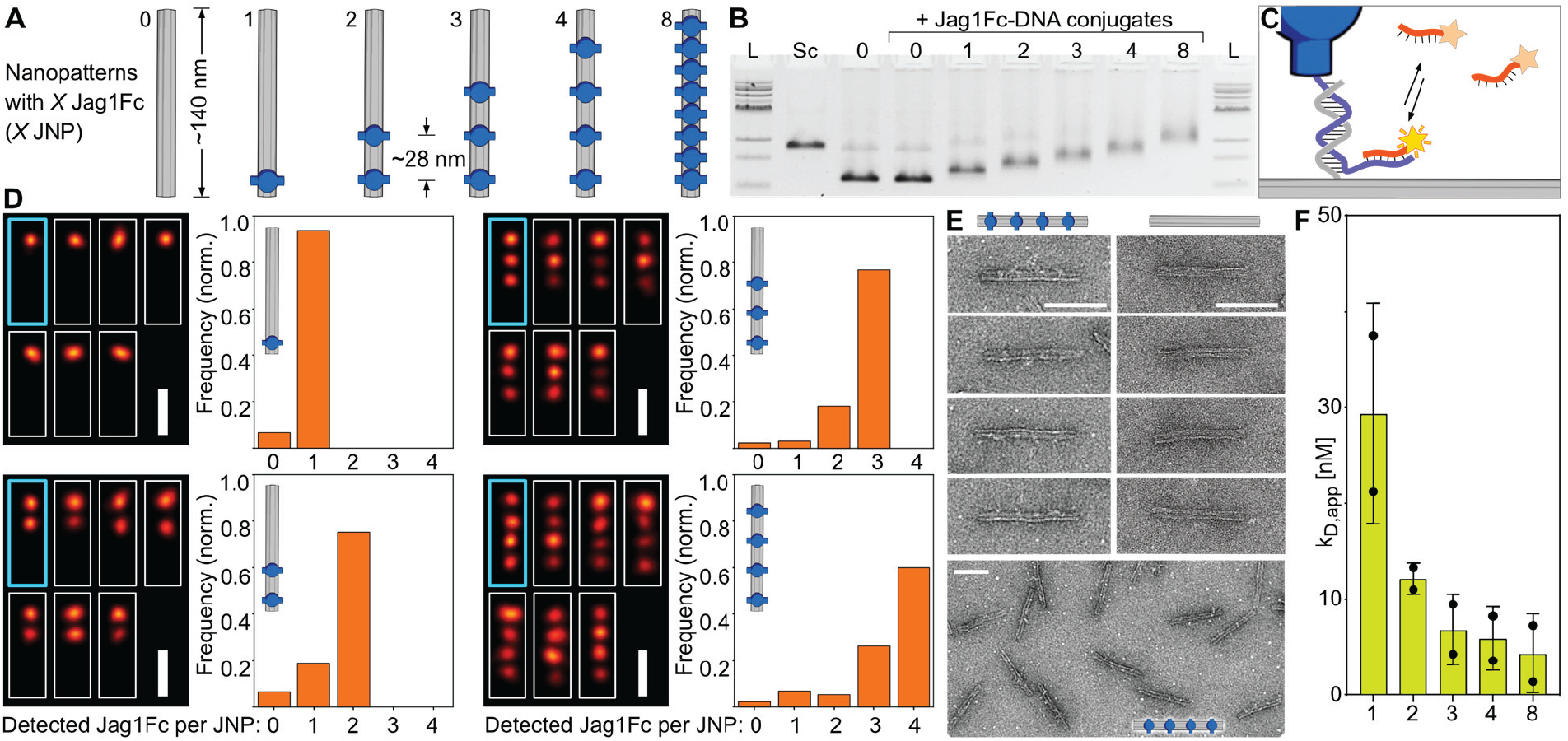
Characterization of Jag1Fc DNA nano-patterns. **(A)** A rod like DNA origami was used to create 1x, 2x, 3x, 4x and 8x Jag1Fc nanopatterns (JNPs). **(B)** Gel retardation assay confirms an increasing molecular weight when increasing the number of proteins positioned on top of the DNA origami. 1Kb ladder (L), scaffold (Sc), 0x JNP (0), repeat of (0) but with added Jag1Fc-DNA conjugates (control for non-specific binding), structures loaded with Jag1Fc patterns: 1x, 2x, 3x, 4x and 8x JNP run on 2% agarose gel stained with ethidium bromide. **(C)** Schematic representation of the DNA origami used for the DNA PATNT experiments, in which the Jag1 proteins conjugated to DNA-handles also contain an extension for DNA-PATNT docking sites used for direct Jag1Fc-DNA conjugate detection (red). **(D)** Average (cyan, thick) and individual cropped DNA PATNT superresolution images (white, thin) of JNPs (scale bars = 50nm) with the bar graphs showing the Jag site occupancy distributions of the different nanorods. **(E)** Zoom-ins of negative stain TEM of (unpurified) 4x JNPs and empty DNA origami rods (0x JNP) and zoom-out of 4x JNPs (bottom). Scale bars are 100 nm. **(F)** Avidity effect of JNPs. Surface Plasmon Resonance measurements using Notch1 (EGF8-12) receptor immobilized on chip the surface and increased concentrations of 1x, 2x, 3x, 4x and 8x JNP flow through the chip. The mean apparent kD of different Jag1Fc nano-patterns is shown on a bar plot where dots represent two individual repeats.

To visualize successful incorporation of proteins on the nanopatterns, we used DNA-PATNT imaging to validate the functionalization states of the different JNPs. For this experiment only the DNA nanostructures were prepared with additional biotin handles for immobilization in the imaging chamber, Cy5 markers for the detection of origami structures with TTRF imaging and Jag1 conjugates carrying DNA-PATNT docking sites for detection of proteins on the structures with DNA PATNT imaging (Figure 2C, Figure S1). For all structures the fraction with the highest detected frequency was the one with the designed number of proteins (1x JNP: 93.5%, 2x JNP: 75.2%, 3x JNP: 76.8%, 4x JNP: 60.0%) with the mean functionalization state being also close to the designed (1x: 0.94, 2x: 1.69, 3x: 2.69, 4x: 3.35) (Figure 2D). Furthermore, the majority of the probes were present as clear monomers (72.6%), with negligible curvature and site-to-site distances closely resembling the designed distances (2x JNP: site 1 to site 2: 26.9±1.7nm; 3x JNP: site 1 to site 2: 27.8±0.9nm, site 2 to site 3: 27.8±0.9nm; 4x JNP: site 1 to site 2: 27.2±1.0nm, site 2 to site 3: 27.2±1.0nm, site 3 to site 4: 27.2±1.0nm) (Figure S1).

We further confirmed the correct geometry of the structures without proteins and the 4x JNPs with negative stain transmission electron microscopy (Figure 2E). The Jag1Fc are barely detectable with TEM probably due to the molecule being on the low molecular weight end for TEM imaging and mostly thin with only the small Fc region being globular. Overall, the gel assays, DNA-PATNT analysis and TEM imaging all confirm that we could produce well controlled monodisperse nanostructures with precise incorporation of Jag1Fc.

To further examine how the ligand multivalency affected binding kinetics in vitro, we used surface plasmon resonance (SPR). We immobilized a biotinylated Notch1 (EGF8-12) protein on a streptavidin modified chip surface and increasing concentrations of JNPs were injected onto the chip. By performing multi-cycle kinetics analysis, we found that the apparent kD of the JNPs decreases when the number of the proteins per nanopattern increases (Figure 2F). Furthermore, the SPR measurements revealed faster association and slower dissociation when more proteins were loaded on the JNPs (Figure S2).

### Jag1 multivalency drive increasing Notch activation

To explore the hypothesis that JNPs can activate and modulate the Notch signaling pathway from solution, we used a model that resembles early neuronal progenitors in human and stimulated these cells from solution with versions of the JNPs (Fig. 3A). Notch activity has been previously reported in iPSc-derived neuroepithelial stem-like (lt-NES) cells where high levels of Notch related genes like *HES1* and *HES5* could be observed^37,38^ while inhibition of the Notch pathway in this system led to neuronal differentiation^37,39^. The model system is motivated by the resemblance of progenitors of the early neural tube, where Notch patterning is believed to be highly relevant for developmental patterning. Tmportantly, the Notch genes in these cells are not genetically engineered to give signal amplification, and should thus provide relevant indications of how an endogenous Notch pathway react to multivalency. Using an antibody against the extracellular part of Notch1, we could confirm the presence of Notch1 using immunostaining in lt-NES cells (Figure 3B) and this was further validated by sequencing data (Figure S3). Moreover, RNA sequencing (RNA-seq) showed that Notch 2, 3 and 4 were also being expressed by these cells (Figure S3). To verify that our experiment was not confounded by a large portion of Notch activation mediated by cell-cell interactions, we initially performed a screen where increasing densities of cells were seeded for 6 hours and stimulated with 4x JNP for 3 hours. We found that a seeding density for JNPs between 9375 and 18750 cells/cm^2^ led to a clearly detectable increase in Notch signaling in response to Jag1Fc DNA nano-patterns (Figure S4A). We further treated iPS cells with 3x JNPs and empty JNPs for 1, 2, 3, 4, 5, 6, 7 hours and measured the expression levels of *HES1* gene and *GAPDH* for each sample, by qPCR. *HES1* expression was normalized to the housekeeping gene *GAPDH*, and the differences in expression levels for 3x JNP to 0x JNP for each time point plotted (Figure 3C). Dynamic expression of Notch target genes is known to be an important effect during developmental processes, particularly in maintenance of neuronal progenitors while stable expression inhibits both processes.^40^ HES1 has been shown to be required for this maintenance, while downregulation of HES1 results in neural differentiation.^41,42^ Another example is the effector gene *Hes7* that oscillates with a 2-hour cycle in presomitic mesoderm (PSM) leading to periodic segmentation of PSM and bilateral pair of somites every 2h.^43^

**Figure 3:**
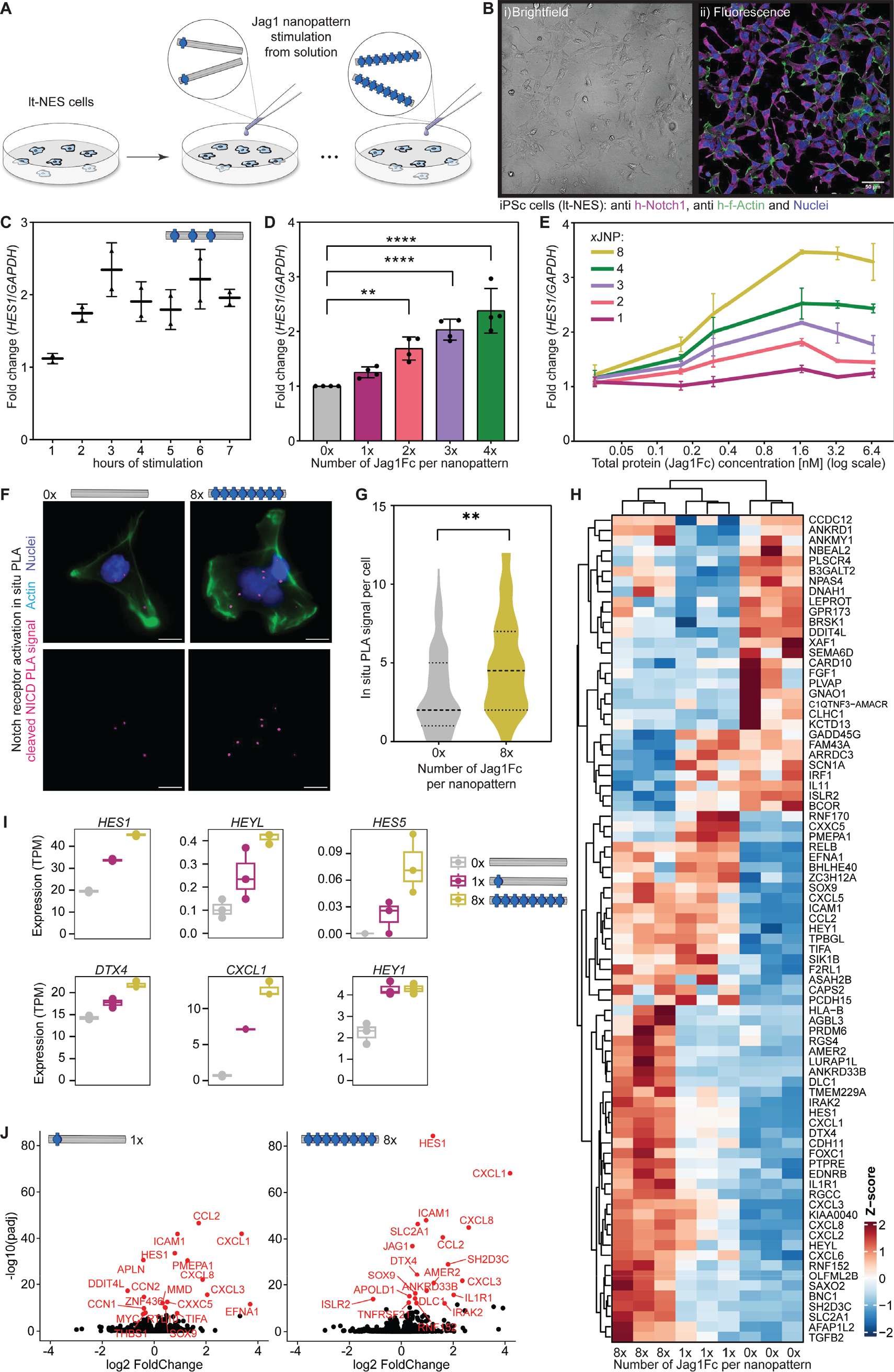
Activation of the Notch pathway by Jag1Fc-nanopatterns. **(A)** DNA nanopatterns (NP) containing 0x, 1x, 2x, 3x, 4x and 8x Jag1Fc (JNPs) were used to stimulate iPSc-derived neuroepithelial stem-like (lt-NES) cells. **(B)** iPS cells shown in i) brightfield and ii) fluorescence channel after immunostaining with antibody against the extracellular part of Notch1 (magenta), F-actin (green) and nucleus (blue). Scale bars, 50 μm **(C)** iPS cells stimulated with 3x JNP for 1, 2, 3, 4, 5, 6 and 7 hours. Relative expression of Hes1 performed with primers for *HES1* and *GAPDH* genes. **(D)** Bar plot of 1x, 2x, 3x and 4x JNP induced activation of Notch pathway detected after RNA extraction and qPCR with primers against *HES1* and *GAPDH* genes. Data points represent 4 biological repeats. One way analysis of variance (ANOVA) was followed by Dunnett multiple-comparison test (**P< 0.01, ****P<0.0001). **(E)** iPS cells stimulated with 1x, 2x, 3x, 4x and 8x JNP at 0.03, 0.16, 0.3, 1.66, 3.33 and 6.66 nM final concentrations. RNA extracted and qPCR assay performed with primers for *HES1* and *GAPDH* genes. **(F)** Proximity Ligation Assay (PLA) performed using antibody against cleaved NTCD on iPS cells treated either with 0x JNP or 8x JNP. Representative images of cells for each condition show with dots the activation signal (magenta), F-actin (green) and nucleus (blue). Scale bars, 10 μm. **(G)** Violin plot of the PLA experiment when detection signal is measured for 50 cells for each condition. One way analysis of variance (ANOVA) was followed by Tukey multiple-comparison test (**P< 0.01). **(H)** Heat map diagram of mRNA sequencing experiment performed on iPS cells after stimulation with 0x, 1x and 8x JNP and three biological repeats for each condition shown for genes with FDR < 0.05 and absolute log2FC > 0.5Data is shown automatically clustered using hierarchical complete-linkage clustering of euclidean distances. **(I)** Depicted Notch pathway related genes and transcripts per million (TPM) plotted for each condition. **(J)** Volcano plotsof genes upregulated by 1x JNP (1) and 8x JNP (8) relative to 0x JNP (0).

From our data we observed that the measured trend of activation levels at different time points appear to follow this known oscillation effect in the *HES1* gene. We could observe a high activation at 3 hours after initiation of JNPs stimulation (Figure 3C), and this condition was then used for the following experiments. Tmportantly, to compare the effect of different JNPs on the Notch activation, we normalized their concentrations to the proteins, and not to the DNA origami itself, assuming a full occupancy of the protruding sites (Note that this method of normalizing, at best leads to an underestimation of total ligands in solution in the samples with multiple sites since these samples tend more to have occasional empty sites, see Fig 2B,D). First, we treated the iPS cells with 3.3nM of total protein on 1x, 2x, 3x and 4x JNP and observed a significant upregulation when more than one protein was placed per nanopattern, with increasing number of Jag proteins per structure resulting in increasing activation (Figure 3D).

To test whether the structures stimulate Notch activation in a dosage dependent manner, we used increasing concentrations of 1x, 2x, 3x, 4x and 8x JNPs, to stimulate iPS cells and followed the receptor activation with real time qPCR measuring products of *HES1* and *GAPDH* genes. We observed that cells stimulated with each structure reached a plateau of stimulation after 1.66nM Jag1Fc, again corroborating that the multivalency appears to be the most important factor (Figure 3E). We also validated that the upregulation of *HES1* was an effect of Notch receptor activation, by adding inhibitors for ADAMs and γ-Secretase respectively 2 hours prior to stimulation, followed by stimulation of the cells with 3x JNPs, observing no *HES1* expression upregulation (Figure S4B). Taken together, these results demonstrate a clear effect of Notch downstream targets being affected by ligand multivalency.

Direct verification of activation of endogenous Notch receptors have been challenging due to a lack of tools available in other receptor systems, for example such as those for tyrosine kinase activated receptors. Here we applied a proximity ligation assay (PLA),^44^ that amplifies the signal from an antibody against a region that is exposed only after γ-secretase cleavage occurs^45^. For this experiment we used a 0x JNP (*i.e*. only the DNA origami rod), to measure the baseline activity, and the 8x JNP which our previous assays had shown to give the strongest stimulation effect. We observed significantly more PLA signal per cell when stimulating with the 8x JNP (Figure 3F, 3G and Figure S4C), verifying that the detected downstream effects were indeed due to Notch being activated.

To further analyze downstream signaling, we stimulated iPS cells with 0x (empty NPs), 1x and 8x JNPs, in three different biological repeats and analyzed the samples with RNA sequencing (RNA-seq). As expected, different genes were upregulated upon stimulating cells with NPs loaded with JAG1FC patterns compared to empty NPs (Figure 3H). Tmportantly, RNA-seq revealed that many Notch-related genes like *HES1, HEYL, HES5, DTX4, CXCL1 and HEY1* were upregulated after stimulation with clusters of Jag1Fc ligands on NPs (Fig 3T). It has been previously described that NTCD forms a complex with DNA binding protein CSL and MAML at the sequence pair sites (SPS) either as monomer or as dimer resulting in different transcription outcomes.^46,47^ Interestingly, we observed that genes dependent on dimerization of NTCD at the transcription complex (NTC) *e.g. HES1* and *HES5*, became upregulated by larger clusters of Jag1, while *HEY1*, which likely depends on monomeric NTCD transcription complex^48^, reached its maximum upregulation already with 1x JNP. The nanopatterns containing 1x Jag1Fc activated the transcriptional response of the Notch pathway, but with a weaker footprint compared to the 8x JNP. Additional visualization using volcano plots showed the highest upregulated genes for 1x and 8x JNPs relative to endogenous levels of genes (0x JNP case), where notably upregulation of the Jag1 gene also suggest successful stimulation of the Notch pathway. (Fig. 3J).

### Multivalency effect persists irrespective of potential force sources

The prevailing hypothesis of the mechanism for Notch receptor activation involves forces generated by the ligand expressing cell when endocytosis of the ligand-receptor complex uncovers the NRR region of the receptor for subsequent cleavage by ADAM metalloproteases^24^. Considering that a repulsion force could be generated between our Jag1 nanopatterns and the cell membrane due to their mutual negative charge (Figure 4A), we shielded the charge of the DNA nanotube by coating it with oligo lysine (K10) solution to reduce the negative surface charge of the probes and thus to reduce any potential repulsive forces between the lipid membrane and the DNA nanostructures. K10 coating has previously been used to neutralize DNA origamis and protect them from low salt denaturation and nuclease mediated degradation.^49^ Screening different K10 concentrations relative to origami concentration we found that at a ratio of 0.5:1 azide to phosphorus groups (N:P) in the DNA, the nanostructures were coated with K10 without forming aggregates which also led to a significantly changed charge as could be seen from gel electrophoresis (Figure S5A). Treating iPS cells with bare and coated Jag1 nanopatterns and comparing their activation effect on Notch pathway, we observed that shielding the negative charge of the DNA nanostructures didn’t reduce the relative *HES1* gene levels, indicating that electrostatic repulsive forces did not mediate Notch activation in our system (Figure 4A).

**Figure 4.**
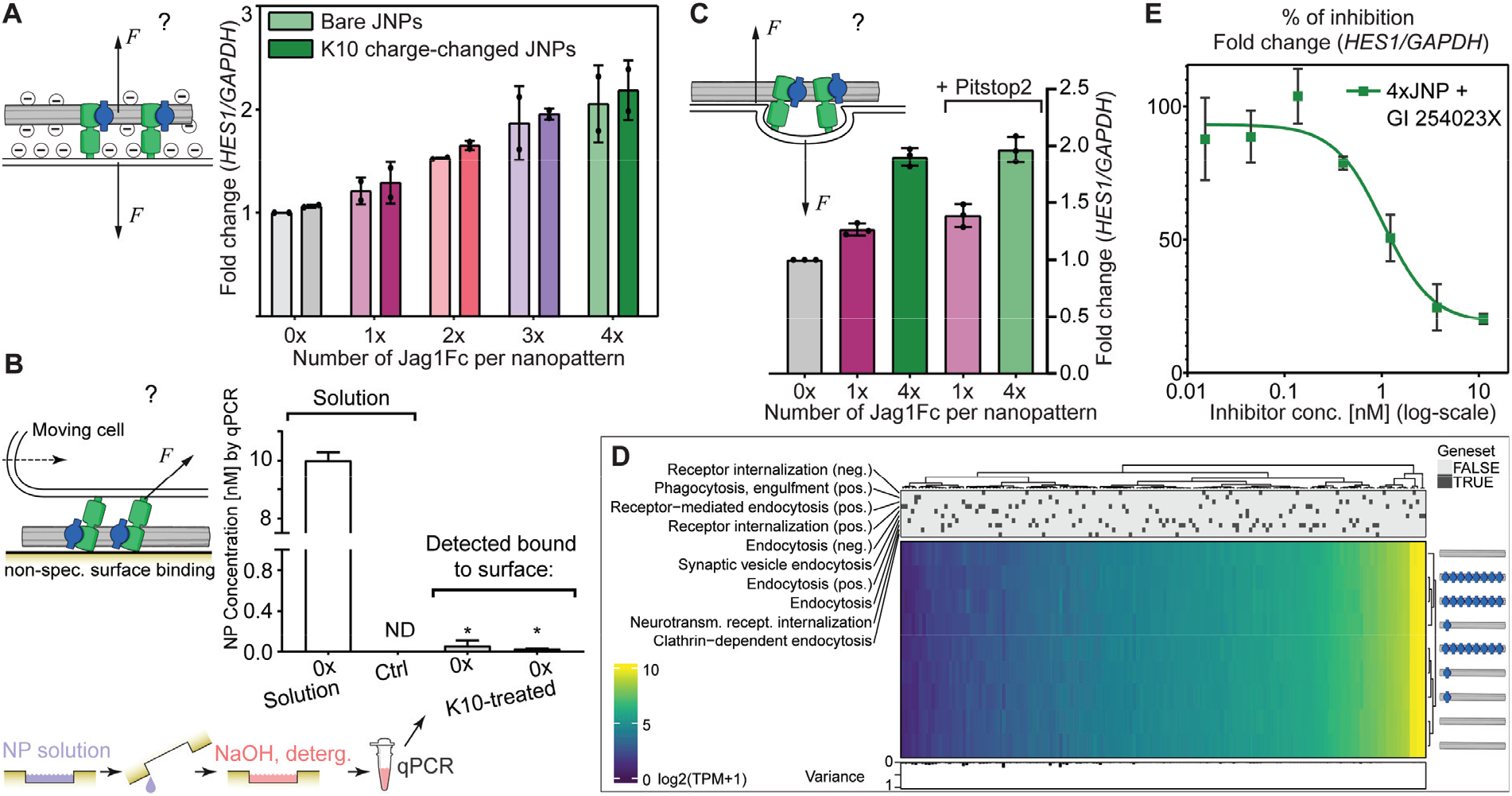
Multivalency drive Notch activation despite potential force sources and ADAM1O is the driving metalloprotease. **(A)** Electrostatic repulsion between negative DNA origami and negative cell membrane could cause pull. By coating the DNA nanopatterns with oligolysine (K10) solution before stimulation their charge is significantly altered (going neutral/positive). TPS cells were stimulated with 0, 1, 2, 3 or 4x JNPs before and after coating with K10. RNA extraction and qPCR was performed with primers against *HES1* and *GAPDH* genes. Dots represent 2 biological repeats. **(B)** Non-specific surface attachment of JNP could drive force generation in motile cells. qPCR measurements directed to part of the DNA origami scaffold (M13mp18 ssDNA) to detect concentrations of structures in solution and, following harsh washing of cell culture surfaces exposed to JNP solution, any remaining non-specifically bound DNA origami. * Denotes measurements below the linear range of the assay. **(C)** Endocytosis could drive force generation. Using Pitstop2, an inhibitor for clathrin-mediated endocytosis on TPS cells stimulated with 1x JNP and 4x JNP followed by qPCR of *HES1* and *GAPDH*, show a similar multivalency effect as the samples without Pitstop2. **(D)** RNA sequencing data of gene ontology genesets that are implied in different levels of endocytosis. Data is shown automatically clustered using hierarchical complete-linkage clustering of euclidean distances. **(E)** Increasing concentrations of ADAM inhibitor GT 254023X added on iPS cells and stimulated with 4x JNP. RNA extracted and qPCR experiment performed with primers for *HES1* and *GAPDH* genes. Error bar lines show 3 technical repeats.

There could further be a force generated by cells moving across structures that are non-specifically bound to the cell culture surface (Figure 4B). To investigate this, we used an assay to probe trace concentrations on surfaces of the DNA origami scaffold using qPCR^50^: We exposed the cell culture surfaces to a solution of typical concentration of JNPs, followed by incubation and then removal of the JNP solution. These cell culture wells were then washed using harsh conditions to extract remnant, potentially non-specifically bound structures, and the solution was subject to qPCR directed to the M13mp18-derived ssDNA scaffold of the DNA origamis. By this quantification, we could observe only traces of DNA origami non-specifically bound, and notably the measurements were below the linear range of the assay (Figure 4B,). The K10 coated origami showed even lower surface attachment. Combined with the results above, this strongly suggests that the multivalency effect we observed in our Notch stimulation, cannot be explained by random surface attachment of JNPs.

It has been hypothesized that one way of generating forces in the Notch signaling system could be from forces generated by invaginations of the cell membrane following initiation of endocytosis of the receptors (Figure 4C). To investigate this, we stimulated TPS cells in the same manner using versions of the JNPs in the presence or absence of an inhibitor against Clathrin-mediated endocytosis. We saw that when the inhibitor Pitstop2 was used, we observed the same multivalency effect as in the stimulation without this inhibitor (Figure 4C). Moreover, investigating expression of the large number of known endocytosis genes in our RNA-seq data, there appeared to be no changes in cells treated with all JNPs tested, 0x, 1x and 8x JNP (Figure 4D). It can be interjected that stimulation may not necessarily trigger transcriptional regulation of these types of basal cellular functions where genes are expected to be reasonably stably expressed, but nanoparticle uptake has nevertheless previously been shown to change transcription of endocytosis genes.^51^ Also, endocytosis inhibitors like the one we use here, are not expected to block endocytosis completely. Nonetheless, the fact that we recorded a multivalency effect in Notch activation independent of endocytosis modulation suggests that the variable strength of effect that follow from the different nanopatterns could at most be very loosely connected to internalization.

The extracellular proteolytic cleavages of Notch receptors are performed by ADAM metalloproteases. Both ADAM10 and ADAM17 have been reported to be able to trigger the release the NTCD but in a context dependent manner. ADAM 10 appears to be responsible for NOTCH activation induced by ligands while ADAM17 is primarily viewed as responsible for ligand independent activation.^13,52^ To assess whether Jag1Fc nanopatterns do induce ligand dependent activation, we used a selective inhibitor for ADAM10 (GT 254023X) in our cell culture experiments. GT 254023X inhibitor has a 100-fold selectivity for ADAM10 over ADAM17. In our experiment, we added increasing concentrations of GT 254023X inhibitor 2 hours prior to cell stimulation with 4x JNPs. The relative inhibition, for samples with increasing concentration of inhibitor, was calculated relative to a sample without added inhibitor (Figure 4E) and the TC50 of this interaction was calculated to be as low as 1,7nM (too low for ADAM17 inhibition) which indicates that ADAM10 is most likely the metalloprotease implied in the activation using JNPs and that the activation is ligand dependent.^52^

### Chimeric patterns suggest a time-of-binding-dependent effect

The question remained whether the multivalency effect we observed is due to clustering of the receptor or if the effect is mainly due to avidity and increased time-of-binding ligand-receptor pairs (Figure 5A). To test whether simply increasing the avidity of the structures without using more Jag1 ligands we designed chimeric structures carrying different moieties that should also bind cell surfaces, placed on the same structure as Jag1Fc.

**Figure 5:**
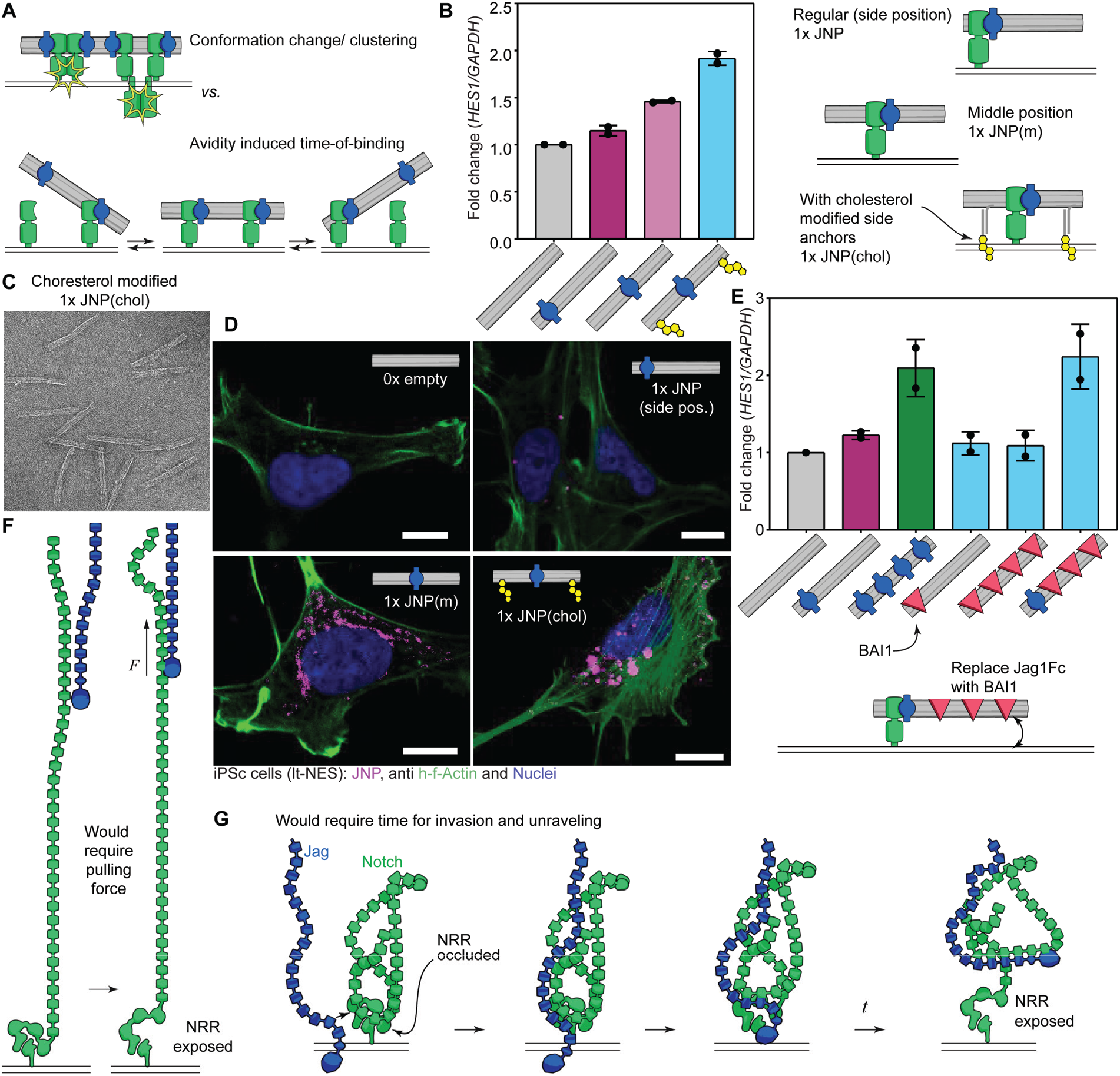
Chimeric structures indicates that multivalency effect is primarily caused by increased binding and suggests hypothesis of tertiary structure occlusion. **(A)** The question arises whether the multivalent patterns of Jag1 induce increased activation due to clustering and potential conformational changes following this, that enhance activation (top). Or, whether the increased multivalency lead to a reduced off-rate of bound structures leading to more time for activation of single receptors (bottom). **(B)** Cholesterol chimeras. Activation data (left) and schematic representation (right) of 1x JNP with protein placed at the side as before, in the middle and in the middle with cholesterol modifications. TPS cells were stimulated with 0x JNP and 1x JNP, 1x JNP(m) and 1x JNP(chol). RNA extracted and qPCR experiment performed with primers for *HES1* and *GAPDH* genes. Symbols represent 2 biological repeats. **(C)** Negative stain transmission electron microscopy of cholesterol modified JNP. **(D)** Confocal microscopy images of iPS cells after stimulation with 0x JNP, 1x JNP, 1x JNP(m) and 1x JNP(chol). NPs contain fluorophores to observe their interactions with cells shown in magenta, actin is shown with green and nucleus with blue. Scale bars are 10 μm. **(E)** BAT1 chimeras - where Jag1Fc is swapped at certain positions with the protein BAT1. Fold change upregulation of *HES1* gene after stimulation of iPS cells with 0x JNP, 1x JNP, 4x JNP, 1x BAT1 NP, 4x BAT1 NP and combination of 1x Jag1Fc and 3x BAT1. Symbols represent 2 biological repeats. **(F)**,**(G)** Different conceptual models of Notch activation. **(F)** Where EGF repeats of Notch (green) extend out like a rod from the cell membrane (lines at the bottom), a force would be able to transport the signal of binding down to the cell membrane to expose the NRR. **(G)** Where EGF repeats of Notch are curled up in a tertiary structure, a binding of ligand would be able to induce unraveling of the tertiary structure over time to allow exposure of the NRR.

First, we positioned a single Jag1Fc in the middle of the DNA nanostructure and added one cholesterol modified DNA strand at each side (Figure 5B). Cholesterol contains a hydrophilic hydroxyl group while the tail of the molecule is hydrophobic known to interact with the lipid membranes.^53^ The structures remained monodisperse as could be observed by TEM of the cholesterol modified structures (Figure 5C). By introducing these cholesterol molecules to the DNA nanostructures, we aimed to increase the interaction time of the JNP with the cell membrane without actually increasing the number of Jag1. As a control, we also produced a similar structure with the Jag1Fc in the middle but without cholesterol anchors. Surprisingly, when Jag1 protein was placed in the middle of the DNA nanostructure it gave a slightly higher activation of the Notch pathway compared to the standard 1x JNP, where the Jag1Fc is placed at the side of the DNA nanostructure (Figure 5B). This effect could possibly be explained by how the horizontal position of DNA origami to the membrane prevents the ligand from being uptaken, which could increase the activation.^54^ Notably though, incorporation of Cholesterol moieties on the JNP caused even higher activation of the Notch pathway, similar to what we previously observed for the 4x JNPs, which suggest to us that a prolonged residence time is the primary factor helping the complex to accomplish a successful interaction. To observe the interaction of these JNPs with the iPS cells we introduced 15 fluorophores (magenta) at the inside of the DNA nanotube structures and after 3 hours of stimulation we fixed the cells, stained for actin (green) and the nucleus (blue), and imaged the sample with confocal microscope (Figure 5D). The results of this experiment showed similarly that when Jag1 was placed in the middle of the NPs with or without cholesterols, it interacted more with the cells compared to 0x JNP and 1x JNP on the side.

To further control for the hypothesis that Notch activation could depend on increasing the time of interaction with the ligands, we produced another chimeric DNA nanopattern in which some of the Jag1 ligands were replaced by BAT1. BAT1 is a protein that interacts with integrin receptors and CD36.^55^ In a previous study co-localization of integrins and Notch1 in neural progenitors indicates either direct interaction of the two proteins or implication of integrins in the Notch intracellular trafficking ^56^. TPS cells were stimulated with chimeric Jag1/BAT1 nanopatterns (Figure 5E and Figure S5B) and we observed that BAT1 protein alone didn’t activate the Notch pathway but when it was placed together with 1x Jag1Fc protein on a DNA nanotube the same activation level was reached as with 4x JNP. We also note that this experiments further rules out charge-effects and molecular weight-effects being important for the multivalency effect we observe, as the BAT1 proteins used here bear similar KpA and molecular weight as the Jag1Fc. Overall, the results of the chimeric structures indicate that the increase in Notch activation with multivalency that we observe, is primarily due to an increase of binding time at the receptor.

## DISCUSSION

Here we have provided evidence that by using nanoscale control of patterns of Jag1, it is possible to activate Notch from solution without exerting pulling forces on the receptor. That a force-dependent mode of activation would also exist is not contradicted by our data - these modes could be complementary and one could be more dominant. Biophysical and cell-biology studies have provided support for the hypothesis that Notch ligands primarily induce activation by cell-cell contacts and force exertion on the receptor.^24–26,28^ Nevertheless, there are also a number of studies suggesting long-range activation (without cell-cell contact) or activation by secreted Notch ligands.^57,58^ As well as novel techniques to stimulate Notch, probably by a mechanism similar to what we see here, using ligand coated beads in solution^59^ or antibody-clustered Notch ligands^60^. How these observations would be explained using exclusively a force dependent mode of activation is less clear.

We argue that support for the current, stretched-rod structural model of Notch, which has been used to exclude other mechanisms than force, is incomplete. Without itself being experimentally verified, the stretched-rod model of the Notch ECD has to some extent provided indirect support for the pulling force model: If indeed, the receptor confirmation is that of a stretched rod (Figure 5F), then it is hard to explain how the crucial region close to the membrane (the Negative Regulatory Region, NRR) would be affected other than via a force signal traveling down the length of the receptor. Because the ligand-binding domain is far from the membrane along the primary sequence (Figure 5F), a pull on the extremity would be required to induce a conformation change on the distant NRR. While the structure is well characterized in terms of the ligand binding domain^21^ and the NRR^11^, a complete picture of the extracellular domain of Notch receptors remains unsolved, and data from pairs of EGFs supports the existence of mostly flexible joints between them^61^. If the EGF repeats act more as beads on a string which appears to be the case, the biophysical concept of an entropic spring would actually favor a condensed-, as opposed to a stretched-, structure, even without intramolecular interactions. One of the few attempts at looking at the entire ECD of Notch with electron microscopy, show bundled up structures whose size would be incompatible with a stretched rod model.^62^

If the Notch ECD is curled up in a tertiary structure like this suggests (Figure 5G), other types of feedback from the binding domain to the NRR could be possible. Our data on activation induced by Jag1 nanopatterns and chimeric patterns favors an explanation where more activation is induced by reducing the off rate of the structures by increased binding to the membrane via either more Jag1, or other molecules (cholesterol, Bai1). This suggests a slower mode of activation that could be another way of exposing the NRR region for ADAM10 cleavage. Similar to DNA strand invasion^63^ (a slow reaction where a random walk of bound bases eventually leads to the release of a bound DNA region) the Jag1 could by its binding, invade a ravel of Notch EGFs that eventually leads to release of material from the NRR and its subsequent exposure (Figure 5G). Note that in this model, a pulling force exerted on the ligand, would probably act as a catalyst for the reaction as the pulling away of bound EGF regions would speed up the exposure of the NRR. In that sense, this hypothesis would stand as a complement to the force model, and not necessarily as a contradiction. This view of a more complex interaction between the Notch ECD and its ligands has been suggested before, after studying deletion mapping and binding,^64^ antibody binding interference^65^ and, more recently, cross-linking mass spectrometry of Notch^66^. But that this type of binding could lead to a slower, force independent mode of activation has not previously been suggested.

Another alternative activation mechanism that is also compatible with our data would be something akin to a kinetic segregation model.^67^ In this model for T cell activation a balance between kinase and phosphatase is disturbed when the bulky CD45 phosphatase is assumed to be excluded from the close synapse forming upon T cell receptor binding, thus toppling the balance in favor of the CD79 kinase which then initiates the signaling. *I.e*. opposite to clustering, a segregation of molecules would lead to activation. It is possible that the presence of the bulk of a 5 MDa DNA origami nanostructure, could be enough to disturb the local balance between ADAM10 and some regulatory counterpart that gets sufficiently excluded under the origami to facilitate the activation. Our results where placing a single Jag1Fc in the middle of a nanostructure, as opposed to one of the ends, lead to slightly higher activation, might be related to this model. That ADAM10 could rely on other regulatory factors has been suggested before^14^ we suggest that these might be inhibitory, and their exclusion required for proper cleavage.

Due to the fact that we are using Jag1Fc and not monomeric Jag1 binding domains, we cannot exclude that dimerization of Notch receptors are important for activation. However, the results concerning multiple Jag1Fc patterns, combined with the results from chimeric structures, point to an effect of multivalency, leading to an increased time of binding, as opposed to an effect of receptor clustering induced by the JNPs (because if the latter is the requirement, why do the chimeric structures increase activity). Future experiments similar to what we present here where a suitable production of monomeric Jag1-DNA conjugates is developed, could provide an answer to whether dimerization is in fact needed as a minimum for activation.

It is interesting to compare our results with studies on synthetic Notch receptors. In SynNotch receptors,^68^ only the transmembrane- and regulatory-(NRR) regions are kept intact, whereas the rest of the Notch receptor, both the ECD and TCD are replaced by artificial binder regions and transcription regulators, respectively. These artificial Notch-like constructs are assumed to require cell-cell contact for activation, which is explained with a force model. Interestingly, a recent study investigating differences between SynNotch and WT Notch found that SynNotch, as opposed to WT Notch, does not require an intracellular domain on the ligand cell side to initiate activation.^69^ Another type of synthetic Notch, called SNTPR, has recently been introduced where the NRR is replaced altogether.^70^ When comparing the activation of these systems with our data on endogenous Notch, a complex picture arises. In SynNotch, the EGF domains are replaced, but the activation is still dependent on ADAM10 and γ-Secretase. Although we did not test SynNotch with our nanopatterns, one could argue that these earlier results, taken together with our new data, appear to favor some type of kinetic segregation model as an explanation, as the EGF-unraveling hypothesis would not apply to these artificial Notch mimics. In particular the SNTPR-constructs, that appear to be both dependent on ADAM10 and γ-Secretase for activation despite sometimes lacking the NRR altogether, would be difficult to explain with a force model (as well as with the EGF-unraveling model).

In this study, we have focused on the low binding affinity Jag1 ligand. Whether or not other ligands could drive Notch activation with similar differences in outcome due to multivalency, remains to be investigated. Ligand specific behavior of the Notch receptor has been observed, even with paralogs of the same family. For example, expression of DLL1 and DLL4 leads to opposing effects in muscle differentiation.^71,72^ Recently, a study showed that DLL1, a low binding affinity ligand, activates Notch1 in pulses by targeting Hes1 genes while DLL4, a higher binding affinity ligand, activates Notch1 in a sustained manner by upregulating Hey1 and HeyL genes.^73^ Similar to what is argued in that work, we propose that lower affinity Notch ligands, like Jag1, would rely more on multivalency to drive Notch activation than high affinity ligands.

In addition to providing insight to the basic mechanisms of Notch activation, the results we present here lay the foundation for an alternative development strategy for new soluble Notch agonists. These are currently an elusive and highly sought-after class of drug. Attenuation of Notch signaling due to tumor growth has been shown to cause immunosuppression that could be overcome by enhancing Notch activation in the hematopoietic microenvironment.^60^

Targeted Notch activation can also be beneficial in several other diseases: it inhibits acute myeloblastic leukemia growth and increases survival,^74^ it is suggested as a treatment for Notch ligand loss of function diseases like Alagille Syndrome and for regenerative medicine.^75,76^

In conclusion, using well-defined molecularly precise patterns of Jag1 ligands we provide evidence that the Notch pathway can be activated from a solution phase in a manner that is not dependent on a force activation mechanism. Instead we show that by increasing the number of Jag1 ligands, we increase the activation efficiency. The fact that this effect remains even when one Jag1Fc is combined with other binders (cholesterol, Bai1) makes it difficult to explain this effect through a clustering model. This leads to two possible conclusions: (i) either increased avidity and the time of binding at the receptor is enough to initiate the pathway, potentially via an unraveling of the EGF domains from the NRR, or (ii) the bulk of the nanostructure together with long enough binding time, mimics a neighboring cell in a way sufficient to modulate a kinetic segregation model of unknown players, most likely related to ADAM10.

## MATERIALS AND METHODS

### Jag1 protein production

Jag1 plasmid was a gift by Susan Lea and Jag1 protein was produced like previously described ^10^. Jag1Fc protein was produced in human embryonic kidney 293T (HEK293T) cells by using transient transfection. Plasmid and transfection reagent (lipofectamine 2000) were mixed at a ratio 1:3 in optimem media and added into cells. After 1 day media collected and replaced with 10% FBS DMEM and cells were let to produce proteins for 3 more days. Proteins containing His tag at the C terminus were purified with affinity purification column His trap FF.

### Jag1 conjugation to DNA

Jag1Fc containing 6x Histidine at the C terminus reacted with the chemical Bis-sulfone-DBCO. The bis-sulfone group reacts with the histidines at the C terminus of the protein, and then an azide modified oligonucleotide was conjugated to the protein by click chemistry between the azide-and the DBCO-group.

### Culture of Neuroepithelial Stem (NES) cells AF22

NES cells were cultured as adherent cells on cell culture flasks previously coated with 20 μg/ml polyornithine (Sigma) for 1 hour and 1 μg/ml Laminin2020 for 4 hours (Sigma). Cells were cultured on NES culture medium contained DMEM/F12+GlutaMax (Gibco), supplemented with 10 μl/ml N-2-supplement (100x, Thermo Fisher Scientific), 10 μl/ ml Penicillin-Streptomycin (10,000 U/ml, Thermo Fisher Scientific), 1 μl/ml B27-supplement (50×, Thermo Fisher Scientific), 10 ng/ml of bFGF (Life Technologies) and 10 ng/ ml of FGF (PeproTech). The culture medium was replaced every second day. The NES cells were passaged enzymatically when reaching 100% confluency using Trypsin-EDTA (0.025%, Thermo Fisher Scientific). NES cells were seeded at a density of 40,000 cells/cm^2^.

### Stimulation of Nes cells with Jag1 nanopatterns

Nes cells were seeded at the density of 18750 cell/cm^2^ and let inside the cell culture incubator to attach for 6 hours. Cells were stimulated with different Jag1Fc nanopatterns for 3 hours when we performed either RNA extraction for real time q-PCR and mRNA sequencing experiment or fixed for proximity ligation and other microscopy experiments.

### Inhibition of Notch pathway

Inhibitors of Notch pathway applied on cells 4 hours after seeding the cells or 2 hours before stimulation with JNPs. In this study we used inhibitor of ADAM10 metalloproteases GT254023X (Sigma), 10μM of γ-secretase inhibitor DAPT (Sigma) or 10μM of a general matrix metalloproteinase, MMPs, inhibitor Batimastat (Sigma).

### RNA extraction

RNA extraction for samples used in RT-qPCR experiments was performed using the Cells-to-CT kit (A25599, Thermo fisher) according to manufactures instructions. In most of the experiments 6000 cells were seeded at a 96 well plate. RNA extraction for samples used in RNA-sequencing experiments was performed by using RNeasy Micro Kit (Qiagen).

### Real time-qPCR

cDNA and RT-qPCR experiments performed according to the instructions included on the Cells-to-CT kit. The pcr reaction mixture includes 10μl of SYBR green, 1 μl of primers (250nM final concentration), 5 μl of water and 4 μl of cDNA. Primers used in this study are *HES1* Fw: AGG CGG ACA TTC TGG AAA TG *HES1* Rev: TCG TTC ATG CAC TCG CTG A *GAPDH* Fw: ACT TCA ACA GCG ACA CCC ACT *GAPDH* Rev: CAC CCT GTT GCT GTA GCC AAA

### Proximity Ligation Assay

After stimulation, cells were fixed with 4% final concentration of methanol-free formaldehyde (Thermo Fisher Scientific, cat. no. 28908) for 15 min at 37°C. The cells were washed at room temperature three times for 5 min each with 1x PBS (Sigma Aldrich, cat. no. 806552). Cells were then permeabilized with 1x PBS 0.1% Triton X-100 (Sigma Aldrich, cat. no. 93443) for 15 min at room temperature and washed at room temperature three times for 5 min each with 1x PBS. The samples were blocked with Duolink Blocking Solution (Sigma Aldrich, cat. no. DUO92002) for 1 hour in a pre-heated humidity chamber at 37°C. The rabbit TgG monoclonal antibody for the detection of cleaved Notch1 (Val1744) (D3B8) (Cell Signalling Technology, cat. no. 4147) was diluted 1:200 in 1x Duolink Antibody Diluent (Sigma Aldrich, cat. no. DUO92002). The cells were incubated with the primary antibody overnight at 4°C and washed at room temperature three times for 5 min each with 1x Duolink *In situ* Wash Buffer A (Sigma Aldrich, cat. no. DUO82049). The Duolink *In Situ* PLA Probes anti-rabbit PLUS (Sigma Aldrich, cat. no. DUO92002) and MTNUS (Sigma Aldrich, cat. no. DUO92005) were then diluted 1:5 in Duolink Antibody Diluent and incubated with the sample in a humidity chamber for 1 hour at 37°C. The probes were washed with 1x Duolink *In Situ* Wash Buffer A three times for 5 min each at room temperature. Next, the Ligase was diluted 1:40 in 1x Ligation Buffer (Sigma Aldrich, cat. no. DUO92008) and incubated in a humidity chamber for 30 min at 37°C. The ligation solution was washed from the cells three times with Wash Buffer A for 5 min each at room temperature. The Polymerase was diluted 1:80 in 1X Amplification Buffer (Sigma Aldrich, cat. no. DUO92008) and amplification of the rolling circle amplification product was carried out in a humidity chamber for 100 min at 37°C. The cells were washed three times for 10 min each at room temperature with 1x Duolink *In situ* Wash Buffer B (Sigma Aldrich, cat. no. DUO82049) followed by two washes of 2 minutes with 0.01x Duolink *In Situ* Wash Buffer B. The F-actin of the cells was fluorescently labelled with Alexa Fluor™ 488 Phalloidin (Thermo Fisher Scientific, cat. no. A12379) according to the manufacturer’s instructions. Nuclei were stained with DAPT solution (Abcam, cat. no. ab228549) diluted to a final concentration of 2 μM in 1x PBS and incubated for 40 min at room temperature. Finally, the cells were washed three times for 5 min each at room temperature with 1x PBS. Cells were imaged in 1x PBS with a Nikon Eclipse Ti-E inverted microscope (Nikon Instruments) using a 1.49 NA CFT Plan Apo TTRF 100x Oil immersion objective (Nikon Instruments). The sample was illuminated at a low angle of 15° using the iLAS2 system (Gataca systems) with lasers listed later (Table 1.) using a custom input beam expansion lens (Cairn). The excitation light was filtered with a filter cube (89901, Chroma Technology), an excitation quadband filter (ZET405/488/561/640x, Chroma Technology) and a quadband dicroic (ZET405/488/561/640bs, Chroma Technology). The emitted light was first filtered with a quadband emission filter (ZET405/488/561/640m, Chroma Technology) and additional respective emission filter described later (Table 1.) (Chroma Technology) and the signal was recorded with an iXon Ultra 888 EMCCD camera (Andor) using the Micromanager software with camera parameters describe later (Table 1.). Each condition was tested with two biological replicates consisting of 50 different cells imaged with z-stacks containing 10 planes with a step size of 4 μm. The z-stacks of the *in situ* PLA signal were converted into Maximal Intensity Projections (MTP) using Fiji. From the z-stacks of the F-actin and the nuclei the focused slice was automatically found using Fiji, the images were then pre-processed by adjusting the contrast and brightness and applying a Gaussian blur to improve object detection by thresholding. The nuclei and boundaries of the cells and PLA signals were identified using batch processing with CellProfiler (www.cellprofiler.org).

**Table 1:** Camera and illumination parameters used for collecting PLA data

| Channel | Laser used | Emission filter used | Electron multiplication | Pixel read out rate | Exposure time |
| --- | --- | --- | --- | --- | --- |
| PLA | OBIS 561nm LS, 150mW | ET595/50m | 50 | 10MHz | 400msec |
| Membrane dye | Omicron LuxX+, 488nm 200mW | ET525/50m | 50 | 10MHz | 50msec |
| DAPI | Omicron LuxX+, 405nm 300mW | ET450/50m | 50 | 10MHz | 50msec |

**Table 2:** DNA PAINT filtering parameters

| **Dataset** | **Localization precision cut-off [nm]** | **Ellipticity cut-off** | **Photon count cut-off [count]** |
| --- | --- | --- | --- |
| **Monomer** | 3,5 | 0,15 | 27894,82632 |
| **Dimer** | 4 | 0,15 | 26230,86006 |
| **Trimer** | 3 | 0,1 | 29549,99792 |
| **Tetramer** | 4 | 0,15 | 27975,19826 |

### Immunostaining of Notch1 receptor

Cells were seeded for 6 hours at a density of 75000 cells/cm^2^ and fixed with 4% methanol-free formaldehyde for 15 minutes. Then cells were permeabilized with 0.1% Triton™ X-100 in PBS for 10 minutes, washed twice in PBS and blocked for unspecific binding of antibodies with 3% BSA for 30 minutes. Cells were probed with NOTCH1 Monoclonal Antibody (Thermo Fisher SCTENTTFTC, # MA5-11961) in 3% BSA at a dilution of 1:100 and incubated overnight at 4 °C in a humidified chamber. The next day, cells were washed with PBS and incubated with 1 μg/mL of Goat anti-Mouse TgG (H+L) Cross-Adsorbed Secondary Antibody, Alexa Fluor™ 488 (Thermo Fisher SCTENTTFTC, #A-11001) at room temperature in the dark for 1 hour. At the last, we washed the cells with PBS and stained F-actin and nucleus with Texas Red™-X Phalloidin (Thermo Fisher SCTENTTFTC, #T7471) and NucBlue™ Fixed Cell ReadyProbes™ Reagent (Thermo Fisher SCTENTTFTC, #R37606), respectively, via following the product protocols. Cells were imaged on Zeiss LSM980-Airy2. Post-processing of images was done with Fiji.

### Confocal imaging of Cy5-labeled DNA nanostructures

Cells seeded and stimulated as described above. Cy5-labeled Jag1 nanopatterns used at 1,66nM final concentration. Cells were fixed with 4% methanol-free formaldehyde for 15 minutes and permeabilized with 0.1% Triton™ X-100 in PBS for 15 minutes. Then cells were washed with PBS and stained for F-actin and nucleus with Alexa Fluor™ 488 Phalloidin (Thermo Fisher SCTENTTFTC, #A12379) and NucBlue™ Fixed Cell ReadyProbes™ Reagent (Thermo Fisher SCTENTTFTC, #R37606), respectively, via following the product protocols. Cells were imaged on Zeiss LSM980-Airy2. Post-processing of images was done with Fiji.

### Design of DNA origami nanostructures

The design of the 18 helix-bundle nanorod was done using caDNAno (https://cadnano.org) using hexagonal lattice design parameters for 3D DNA origami described in ref ^33^. The origami design follows the one used in ref ^34^ and has been uploaded to a repository (https://nanobase.org) where the design files and structure files can be downloaded. By folding the structure either with 5’ protruding staples, or empty site staples, respectively, placement of DNA-protein conjugates can be done at will along the top edge of the structure as described in ref ^34^.

### Folding DNA nanostructures

The standard folding conditions used in this study were as follows: 20 nM ssDNA scaffold, 100 nM per staple, 13 mM MgCl2, 5 mM Tris, pH 8.5. Single stranded DNA that serves as scaffold for DNA nanostructures folding was produced, extracted and purified from M13 phage variant p7560 cultured by inoculating E. coli JM109. The approximately 200 single stranded DNA oligonucleotides helper strands (staples), were purchased from Integrated DNA Technologies. Folding was carried out by annealing at 65°C for 4min, then 65 °C to 50 °C for 1min/0.7 °C, 50 °C to 35 °C for 1h/1 °C and 20 °C forever until retrieved. Removal of excess staples was done by washing (repetitive concentration and dilution for seven times) with PBS, pH 7.4, 10 mM MgCl2 in 100-kDa MWCO 0.5-ml Amicon centrifugal filters (Millipore). Samples were diluted to 450 μl and transferred to a prewetted centrifugal filter and centrifuged at 10,000*g*, for 1 min, and then diluted again to 450 μl, mixed well and centrifuged again under the same conditions. Sample was collected via inverting the filter on an Eppendorf tube and centrifugation at 1,000*g* for 2 min.

### Jag1Fc nanopatterns

The ligand Jag1 conjugates were added with a twenty times excess to each protruding site on the DNA nanostructure and incubated in the PCR machine with a temperature ramp starting from 1 h at 37 °C followed by 14 h at 22 °C, and immediately after incubation the nanopatterns were stored at 4 °C. Removal of Jag1 conjugates in excess performed in FPLC system and a size exclusion purification column (Superose 6 Increase 10/300GL, Cytiva). After purification fractions of the peak corresponding to Jag1 nanopatterns collected and concentrated in 30-kDa MWCO 2-ml Amicon centrifugal filters (Millipore). Sample concentration estimated with agarose gel electrophoresis by loading sample of known concentration before purification and sample after purification of unknown concentration. By comparing the intensity of the bands we calculate the final concentration of each sample.

### Agarose gels for characterization of the Jag1 nanopatterns

We prepared 2% agarose gels with 0.5x TBE supplemented with 11 mM MgCl2 (Sigma-Aldrich) and 0.5 mg/ml ethidium bromide (Sigma-Aldrich). We typically loaded 4 μl of 20 nM DNA nanostructures in each lane and ran the gels in 0.5x TBE with 10 mM MgCl2 at 90 V for 3 h, cooled in an ice-water bath. The gels were imaged on a GE Tmage Quant LAS 4000 system.

### Surface plasmon resonance (SPR)

A BTAcore T200 (GE Healthcare) was used to measure the binding kinetics of Jag1Fc nanopatterns to Notch receptor. Streptavidin (Sigma-Aldrich) was dissolved in 100 mM sodium acetate buffer, pH 4.5, and immobilized on a CM3 chip (GE Healthcare) according to the manufacturer’s instructions. The biotinylated extracellular domain of the human Notch1 receptor (8-12) was immobilized at 200 RU. Jag1Fc DNA nanopattern samples were diluted to concentrations ranging from 2,5 nM to 10 nM in PBS, pH 7.4, supplemented with 10 mM MgCl2. The flow rate of the samples was adjusted to 5 μl/min, and a total amount of 35 μl was injected. Sensorgram data were processed with BTAevaluation 3.2 software (GE Healthcare).

### Transmission electron microscopy (TEM)

We applied 3 μl of the DNA nanostructures on glow-discharged, carbon-coated Formvar grids (Electron Microscopy Sciences), incubated for 20 s, blotted off with filter paper, and then stained with 2% (w/v) aqueous solution of uranyl formate supplemented with 20 mM NaOH. followed by a final blot with filter paper. The negative stained samples were imaged by Talos 120V microscope at x and x magnification.

### Sample preparation for DNA PAINT imaging experiments

Versions of Jag1 nanopatterns (JNPs) with one, two, three and four Jag functionalization sites used in other experiments were produced carrying biotinylated anchoring-sites on the opposing side to the Jag sites, along with six internal Cy5 modified staple-oligonucleotides for DNA PATNT independent detection of the nanopatterns. Nanopatterns were produced and purified as described earlier using Jag1 proteins conjugated to oligos containing the anchoring sequence and two DNA PATNT docking sequences. Microscope slides (VWR) and coverslips (1.5H, VWR) were cleaned with acetone and isopropanol and flow-chambers were produced by placing two strips of double-sided scotch tape approximately 0.8cm away from each other on the slides and placing the cleaned coverslips on top of the strips. The channel was first incubated with biotinylated-BSA (Sigma Aldrich)) solution (1mg/mL biotinylated-BSA in Buffer A (10mM Tris-HCl, 100mM NaCl, 0.05% Tween-20, pH 7.5)) for 2 min and was washed with Buffer A. The channel was then incubated with streptavidine (Thermo Scientific) solution (0.5mg/mL streptavidin in Buffer A) for 2 min. Following a washing step with Buffer A the channel was washed with Buffer B (5mM Tris-HCl, 10mM MgCl2, 1mM EDTA, 0.05% Tween-20, pH 8). The channel was then incubated with Jag-nanopatterns solution (50pM Jag-NC in Buffer B) for 5 min followed by washing the channel with Buffer B. Finally, the imager-solution (10nM Atto550-labelled imager strand in Buffer B+ (Buffer B, oxygen scavenger system (2.4mM PCA (Sigma Aldrich) and 10nM PCD (Sigma Aldrich)) and 1mM Trolox (Sigma Aldrich)) was introduced into the channel and the channel was then sealed with epoxy glue.

### DNA PAINT imaging of Jag-functionalized nanopatterns

The DNA PATNT imaging experiments were conducted with a Nikon Eclipse Ti-E inverted microscope (Nikon Instruments) using a 1.49 NA CFT Plan Apo TTRF 100x Oil immersion objective (Nikon Instruments) and a 1.5x auxiliary Optovar magnification. The TTRF illumination was produced using an iLAS2 system (Gataca systems) with an OBTS 561nm LS 150mW laser (Coherent) and an Omicron LuxX+ 642nm 140mW laser (Coherent) and a custom input beam expansion lens (Cairn). The excitation light was filtered with a filter cube (89901, Chroma Technology), an excitation quadband filter (ZET405/488/561/640x, Chroma Technology) and a quadband dicroic (ZET405/488/561/640bs, Chroma Technology). The emitted light was first filtered with a quadband emission filter (ZET405/488/561/640m, Chroma Technology) and an additional emission filter (ET595/50m, Chroma Technology; ET655lp, Chroma Technology) and the signal was recorded with an iXon Ultra 888 EMCCD camera (Andor) using the Micromanager software. For the recording of the Cy5 signal snapshots were taken with 1 second exposure time, 10MHz readout rate and no EM. For the recording of the DNA PATNT data 12000 frames were collected in frame-transfer mode with 300msec exposure time, 10MHz readout rate and no EM gain.

### Quantification of Jag proteins on nanopatterns using DNA PAINT

#### Fitting of localizations

The Picasso software package^77^ was used for preprocessing the raw DNA PATNT data. The Picasso Localize software was used to detect and fit localizations in the raw DNA PATNT movies using the MLE algorithm (Box sixe:7, Min. Net Gradient: 2000, EM Gain: 2, Baseline: 43.2, Sensitivity: 4.1, Quantum efficiency: 0.98, Pixel size: 87nm).

#### Drift correction and filtering of DNA PAINT data

Following fitting the Picasso Render software was used to drift correct the localizations using the redundant cross-correlation (RCC) algorithm (segment size: 200 frames). The data was subsequently filtered using a custom Python script through the removal of low precision localizations and multi-event localizations. The filtered localizations were drift corrected again using the Picasso Render software: individual origami structures were picked using the pick similar tool (pick area diameter:1.5 camera pixel, pick similar std: 1.6), initiated with 20 manually picked origami structures, and the data was undrifted with the undrift from picked feature of the software.

#### Detection of DNA origami

Detection of DNA origamis were performed with a custom Python script. The Cy5 image together with the DNA-PATNT data was used to determine the position of individual DNA origami probes. First the Cy5 image was intensity normalized and a binary image was produced with adaptive thresholding. After noise removal the image was segmented, and the contours were detected for the individual segmented objects. The DNA PATNT data was used in parallel to generate a low-resolution image from where the positions of DNA origami probes were determined by generating a pixel inflated binary image and detecting the contours of objects in a selected size range. The final positions of origami probes were generated from the two set of contour coordinates (Cy5 and PATNT) by combining them: in the case of Cy5 contours with overlapping DNA PATNT contours (distance between contour centers smaller than 90% of sum of contour radiuses) the contour coordinates generated from the PATNT data were used, in the case of Cy5 contours with no overlap (structures without detected Jag1 proteins) the Cy5 contour coordinates were used. Using the center coordinates of these contours and the average size of DNA PATNT contours coordinates for the origami region of interests (ROTs) were determined and DNA PATNT localizations were grouped into these. (Figure S1B).

#### Quantification of Jag proteins within DNA origami ROIs

Reference values used in the later processing steps for neighboring position-to-position distances and linearity scores were calculated using the custom Python script. For the calculation of the position-to-position distances the localizations grouped into individual origami ROTs in the 2xJNP dataset were clustered using the DBSCAN algorithm and the distance between the mean position of the two clusters with the highest number of points in them was extracted. The reference value for the position-to-position distance was then calculated as the mean of the gaussian fit of the resulting distance distribution. (Figure S1C).) For calculating the linearity score cut-off value, the localizations in individual ROTs in the 4xJNP dataset were rendered into a high-resolution image with intensity normalization to the highest pixel value. Localization density maxima were detected as protein positions and after removing outlier points the protein positions were annotated by using their distance matrix. Position to neighbouring position (PTNP) vectors were calculated between the adjacent protein positions and the mean standard deviation of the normal PTNP vectors’ x, y coordinates was calculated as the linearity score for each probe. The cut-off value was then determined as the inflection point of the cumulative distribution of this linearity score for origami ROTs with four detected points (Figure S1D).

Quantification of Jag proteins on individual origami probes residing in the origami ROTs was performed using a custom Python script. Localizations in individual origami ROTs were rendered into a high-resolution image with intensity normalization to the highest pixel value. Initial guesses for the protein positions were determined by detecting localization density maxima in the image and annotating them using their distance matrix after the removal of outliers. ROTs with higher linearity score as the cut-off value and/or with more detected positions than the designed were discarded. In ROTs passing this filtering the initial guess for Jag1 position one along with the position-to-position reference distance was used to calculate the putative regions for each position and these regions were then scanned for local localization maxima to detect density maxima with lower number of localizations. For positions with detected maxima the coordinates of these were then used as the final positions of Jag1 proteins. (Figure S1E).

### Libraries for sequencing experiment

Libraries for RNA sequencing were prepared with TruSeq stranded mRNA (Tllumina 20020595) according to manufactures instructions. Libraries were sequenced on an in-house NextSeq550 using high output v2 kits.

### RNA-seq analysis

Raw data was adapter trimmed and human transcriptome (GRCh38.p13 Gencode v38 protein-coding transcripts; gencode.v38.pc_transcripts) was quantified using Salmon^78^ (v1.3.0, options: −l TSR --validateMappings). Quantifications were summarized to gene-level using tximport^79^ (v1.12.3) and differential expression was calculated as Wald tests using DESeq2^80^ (v1.24.0). Gene ontology enrichment was performed as Fisher overrepresentation tests using PANTHER^81^ (release 20210224, GO database 10.5281/zenodo.5080993 and release 20221103, GO:0030100 Regulation of Endocytosis).

### Statistical analysis

For multiple comparison analysis in Figure 3D, we performed one-way ANOVA followed by Dunnett multiple comparison. Using Prism software (GraphPad), we performed the Shapiro-Wilk normality test, which showed that each group of data followed a normal distribution. For the *in situ* PLA statistical analysis of the processed images (Figure 3F and Figure S4C), one-way ANOVA followed by Tukey post hoc test was performed for multiple-comparison analysis of the four populations comprising the two biological repeats for each condition. Analysis was conducted in a blinded format whereby knowledge of which conditions applied to which groups of cells held by one author was withheld from the author responsible for image acquisition and analysis to avoid bias. After visual inspection of the batch processing, cells that were partially out of the field of view (i.e. cut off) were not included in the analysis.

| **Tukey's multiple comparisons test** | **Mean Diff.** | **95,00% CI of diff.** | **Summary** | **Adjusted P Value** |
| --- | --- | --- | --- | --- |
| 0x JNP (1) vs. 0x JNP (2) | -0,4420 | -1,591 to 0,7066 | ns | 0,7535 |
| 0x JNP (1) vs. 8x JNP (1) | -1,649 | -2,777 to -0,5210 | ** | 0,0011 |
| 0x JNP (1) vs. 8x JNP (2) | -1,618 | -2,711 to -0,5243 | *** | 0,0009 |
| 0x JNP (2) vs. 8x JNP (1) | -1,207 | -2,371 to -0,04367 | * | 0,0386 |
| 0x JNP (2) vs. 8x JNP (2) | -1,176 | -2,306 to -0,04582 | * | 0,0378 |
| 8x JNP (1) vs. 8x JNP (2) | 0,03137 | -1,078 to 1,140 | ns | 0,9999 |

## DATA AND CODE AVAILABILITY

The detailed DNA origami design schematic has been deposited at https://nanobase.org (accenssion 192, https://nanobase.org/structure/192)

Raw sequencing data have been deposited at ArrayExpress (accession E-MTAB-12439) and are publicly available as of the date of publication.

Computational code for RNA-seq analyses and DNA-PAINT image analysis is available at https://github.com/bhogberg/nanoscale-notch-multivalency

## ACKNOWLEDGEMENTS

The authors would like to acknowledge support from the Knut and Alice Wallenberg Foundation for B.H. (Grants KAW 2017.0114 and KAW 2017.0276), from the European Research Council ERC for B.H. (Acronym: Cell Track GA No. 724872), and from the Swedish research council for B.H. (grant no. 2019-01474).

## Author contributions

Conceptualization IS, AIT and BH; Methodology IS and BH; Investigation IS, FF, IRL and YW; Formal Analysis IS, FF, IRL and AL; Resources VCL; Funding acquisition BH; Supervision BH, AIT, BR; Visualization IS, IRL, YW, AL, FF, BH; Writing - original draft IS, BH; Writing - review & editing IS, FF, VCL, BR, AIT and BH

## Declaration of interest

The authors declare no competing interests.

## Notes

### Competing Interest Statement

The authors have declared no competing interest.

