## Supplemental figures for "Notch engagement by Jag1 nanoscale clusters indicates a force-independent mode of activation"

### Clustering effect of Jag1 ligand on Notch signaling pathway

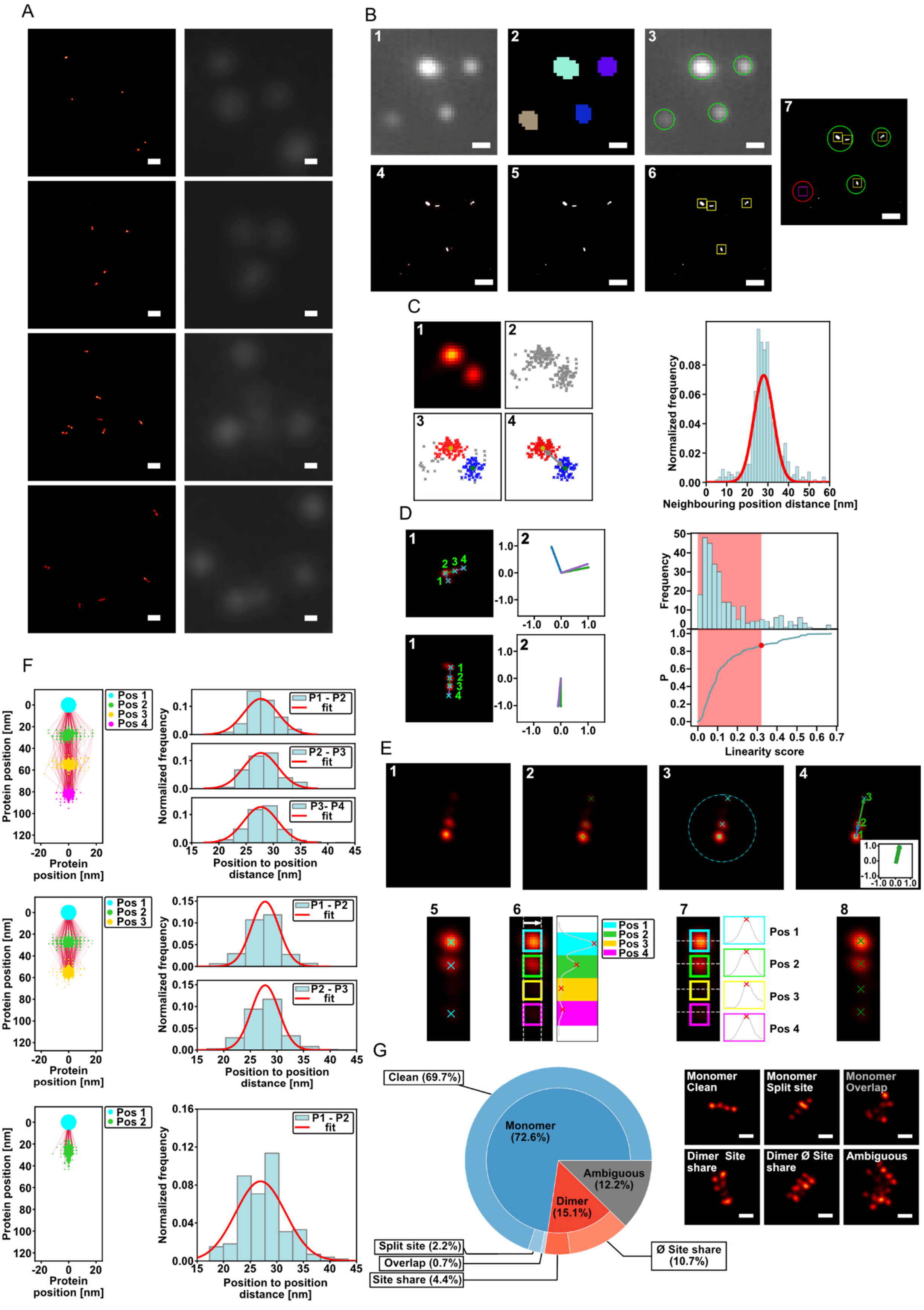
A
B
1
2
3
4
5
6
7
C
1
2
3
4
Normalized frequency
0.00
0.02
0.04
0.06
0.08
0 10 20 30 40 50 60
Neighbouring position distance [nm]
D
1
2
3
4
1.0
0.0
-1.0
-1.0 0.0 1.0
Frequency
50
40
30
20
10
0
P
1.0
0.8
0.6
0.4
0.2
0.0
0.0 0.1 0.2 0.3 0.4 0.5 0.6 0.7
Linearity score
E
1
2
3
4
5
6
7
8
Pos 1
Pos 2
Pos 3
Pos 4
F
Protein position [nm]
Protein position [nm]
Normalized frequency
P1 - P2
P2 - P3
P3- P4
fit
Position to position distance [nm]
15 20 25 30 35 40 45
G
Clean (69.7%)
Monomer (72.6%)
Ambiguous (12.2%)
Dimer (15.1%)
Split site (2.2%)
Overlap (0.7%)
Site share (4.4%)
Ø Site share (10.7%)
Monomer Clean
Monomer Split site
Monomer Overlap
Dimer Site share
Dimer Ø Site share
Ambiguous

**Figure S1: Characterization of Jag1 nanopatterns using DNA PAINT imaging**. **A** Images of 1xJNP (top), 2xJNP (second from top), 3xJNP (second from bottom) and 4xJNP samples (bottom) produced using the DNA-PAINT docking strands on the Jag1 conjugates (left) and the Cy5 origami markers (right) (scale bar = 200nm). **B Steps for DNA origami probe detection:** The intensity in Cy5 raw images was normalized (1), images were thresholded and segmented (2). The probe ROIs (green circles) in the Cy5 images were identified using the contours detection. (3). Low resolution images were generated by rendering the localizations in the respective DNA PAINT datasets (4). A binary thresholded, dilated images were generated, and all contours were detected in them (red circles) (5). Origami ROIs (yellow squares) were identified from contours based on contour area (6). Origami ROIs (yellow squares) with an overlap with a Cy5 ROI (green circles) were retained, for Cy5 ROIs with no overlapping origami ROIs (red circle) origami ROIs positioned around the Cy5 ROIs were generated as empty origami ROIs (magenta square) (scale bars = 500nm) (7). **C Steps for neighbouring position distance reference value calculation** from 2xJNP DNA PAINT dataset (left): High resolution image rendered from localizations lying in the origami ROI (1). Scatter plot of localizations lying in the origami ROI (2). Clustering of localizations in origami ROI using DBSCAN algorithm (red: site 1, blue: site 2, grey: noise) and determination of the center coordinates of positions (yellow: pos 1 center, green: pos 2 center) (3). Calculation of neighbouring position distance using the center coordinates of positions (4). Plot showing the distribution of measured neighbouring position distances (right). **D Determination of origami linearity score thresholding value** from the 4xJNP DNA-PAINT dataset. Origami linearity score calculation presented for a non-linear nanopattern (top left) and a linear nanopattern (bottom left): High resolution image of origami ROI was generated from localization and Jag1 positions were detected (cyan crosses) (1). Positions were annotated (green indices) using pairwise distances and were used to calculate position to neighbouring position (PTNP) vectors (colored arrows) (1). PTNP normal vectors were then determined to be used for calculating the linearity score (mean of the standard deviation of the PTNP normal vectors' x and y coordinates) (2). Plot showing the distribution of calculated origami linearity scores (top right) and the cumulative probability plot (bottom right) calculated from the distribution to determine the inflection point (red spot) used as cut-off value. **E Steps for protein detection and quantification in Jag1 nanopatterns:** Localizations sorted into individual origami ROIs are rendered into a high-resolution images (1). Initial guesses for protein positions are generated (green crosses) (2). Points farther from the center of the ROI window (cyan circle) are excluded to remove random noise (3). Protein positions are annotated (site 1 - site 4) using pairwise distances and the PTNP vectors are calculated to check linearity (4). Images of the origami probes are aligned and cropped using the annotated protein positions (cyan crosses) (5). Initial areas of Jag1 positions (colored rectangles) are generated and the final y coordinates of the protein positions are determined by calculating the sum y intensity profile of the cropped image between the site boundaries and detecting peaks (red crosses) in the resulting plot lying in the respective regions of positions (colored regions) (6). Final x coordinates of the protein positions are determined by detecting peaks (red crosses) in the x intensity profiles generated in the previously determined protein y coordinates (7). Final protein positions are generated (green crosses) using the previously determined x,y coordinates of protein positions (8). **F Characterization of Jag1 nanopattern shapes using DNA PAINT imaging:** Plots showing the positions of Jag1 proteins in aligned individual nanopatterns calculated from the DNA PAINT data (left) with point sizes scaled with point density and with structure trajectories highlighted by lines connecting the positions in individual nanopatterns (left) and histograms showing the Jag1 position to neighbouring position distance distributions (right) for the 4xJNP (top), 3xJNP (middle) and 2xJNP probes (bottom). **G Characterization of Jag1 probe multimerization using DNA PAINT** data generated of the 4xJNP sample: Plot showing the relative frequency of different instances of monomeric and multimeric probes detected in the DNA PAINT data (top) and cropped DNA PAINT images illustrating the different classes (scale bar = 50nm) (bottom).

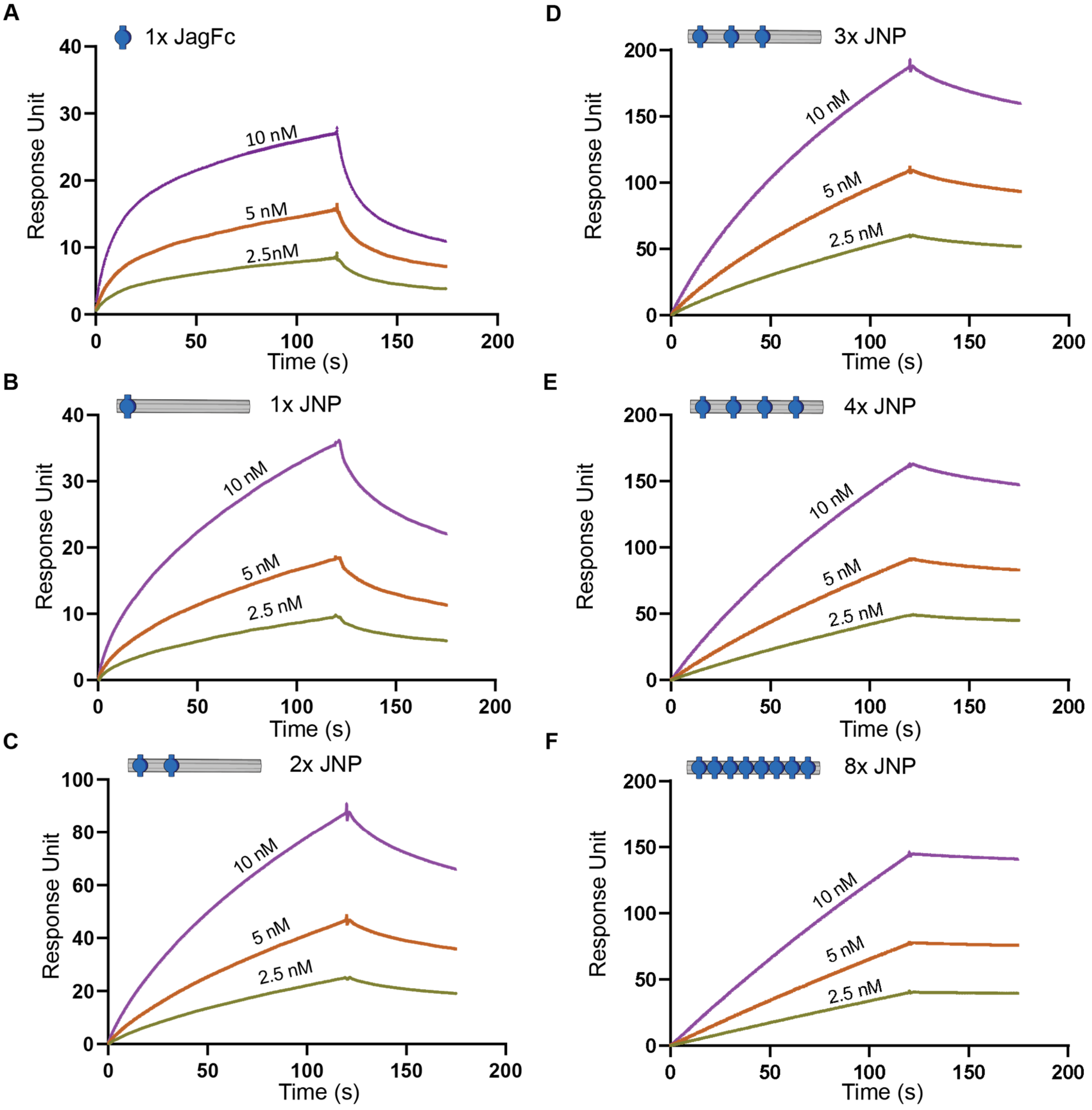


**Figure S2: Binding abilities of Jag1 nanopatterns with Notch receptor measured by surface plasmon resonance.** A short version of Notch receptor (8-12) was immobilized on a CM3 chip surface and increasing concentrations of **A** Jag1Fc **B** 1x JNP **C** 2x JNP **D** 3x JNP **E** 4x JNP and **F** 8x JNP used to perform multi-cycle kinetics on a T200 surface plasmon resonance instrument.

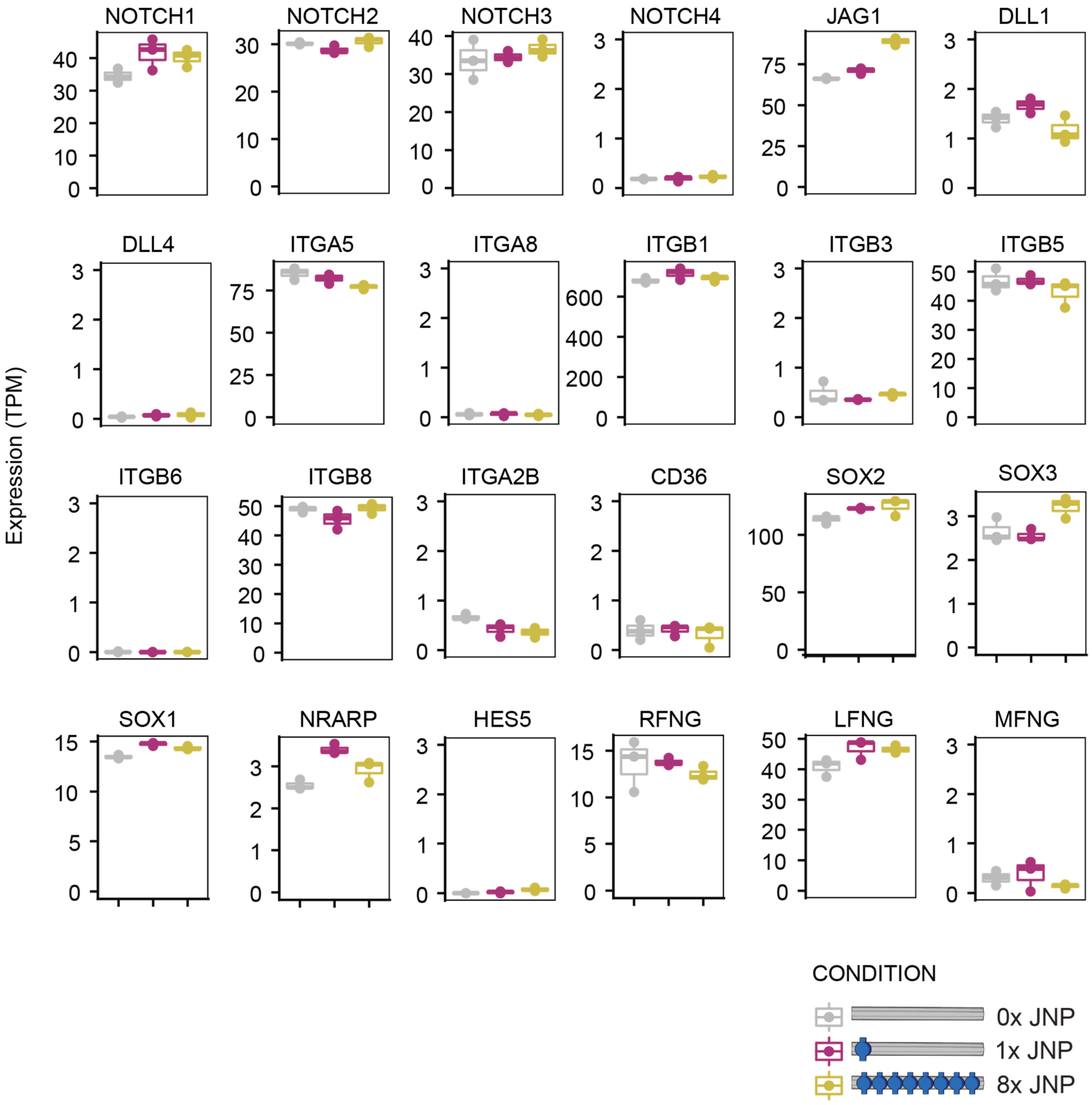


**Figure S3: RNA transcriptomic analysis** of Notch receptor genes (NOTCH1, NOTCH2, NOTCH3, NOTCH4), their ligands (JAG1, DLL1, DLL4), integrin receptors' genes (ITGA5, ITGA8, ITGB1, ITGB3, ITGB5, ITGB6, ITGB8, ITGA2B), CD36, stem cells markers (SOX1, SOX2, SOX3) as well as other Notch related genes (NRARP, HES5, RFNG, LFNG, MFNG). IPS cells were stimulated with 0x JNP (resembles base line levels), 1x JNP and 8x JNP and RNA extracted for RNA sequencing analysis.

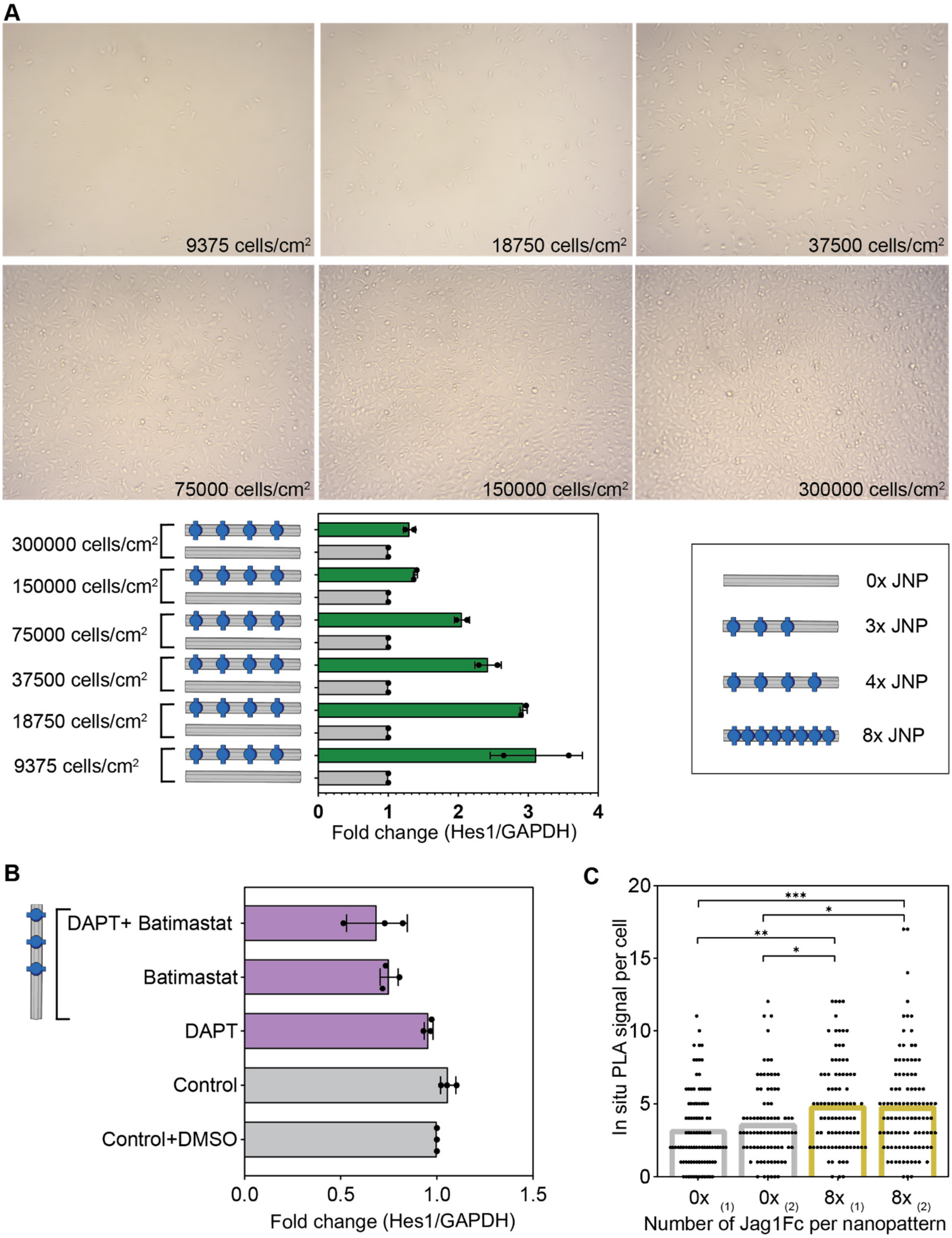
A
9375 cells/cm2
18750 cells/cm2
37500 cells/cm2
75000 cells/cm2
150000 cells/cm2
300000 cells/cm2
300000 cells/cm2
150000 cells/cm2
75000 cells/cm2
37500 cells/cm2
18750 cells/cm2
9375 cells/cm2
0
1
2
3
4
Fold change (Hes1/GAPDH)
0x JNP
3x JNP
4x JNP
8x JNP
B
DAPT+ Batimastat
Batimastat
DAPT
Control
Control+DMSO
0.0
0.5
1.0
1.5
Fold change (Hes1/GAPDH)
C
In situ PLA signal per cell
20
15
10
5
0
***
*
**
*
0x (1)
0x (2)
8x (1)
8x (2)
Number of Jag1Fc per nanopattern

**Figure S4: Stimulation effects of JNPs in iPS cells**. **A Effect of cell seeding density** on the activation of Notch signaling pathway. IPS cells were seeded in 9375, 18750, 37500, 75000, 150000 and 300000 cells/cm$^2$ for 6 hours and stimulated with 0x JNP and 4x JNP. RNA extracted and qPCR experiment performed with primers for HES1 and GAPDH genes. Bar plot shows the mean values and the dots from 2 biological repeats. **B Stimulation of Notch signaling pathway in the presence of Notch inhibitors.** Prior to stimulation metalloproteases inhibitor (Batimastat), γ-secretase inhibitor (DAPT) or only DMSO added to the cells for 2 hours. Then cells stimulated with 0x JNP (control), 0x JNP on the well already containing DMSO (control + DMSO) and 3x JNP on the wells containing the inhibitors. 3x JNP failed to stimulate the pathway when different Notch inhibitors were present on the cell cultures. Bar plot shows the mean values and the dots represent 3 technical repeats of the same experiment. **C Proximity ligation assay detects activated Notch NICD.** Two individual experiments performed when iPS cells stimulated with 0x JNP and 8x JNP shown on the plot. Bar plots indicate the mean in situ PLA signal and black dots the number of dots per cell as measured in 50 cells per condition. One way analysis of variance (ANOVA) was followed by Tukey multiple-comparison test (*P< 0.1, **P< 0.01, ***P<0.001).

**A**

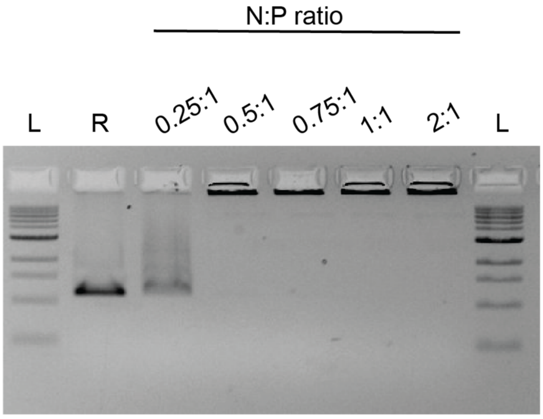


**B**

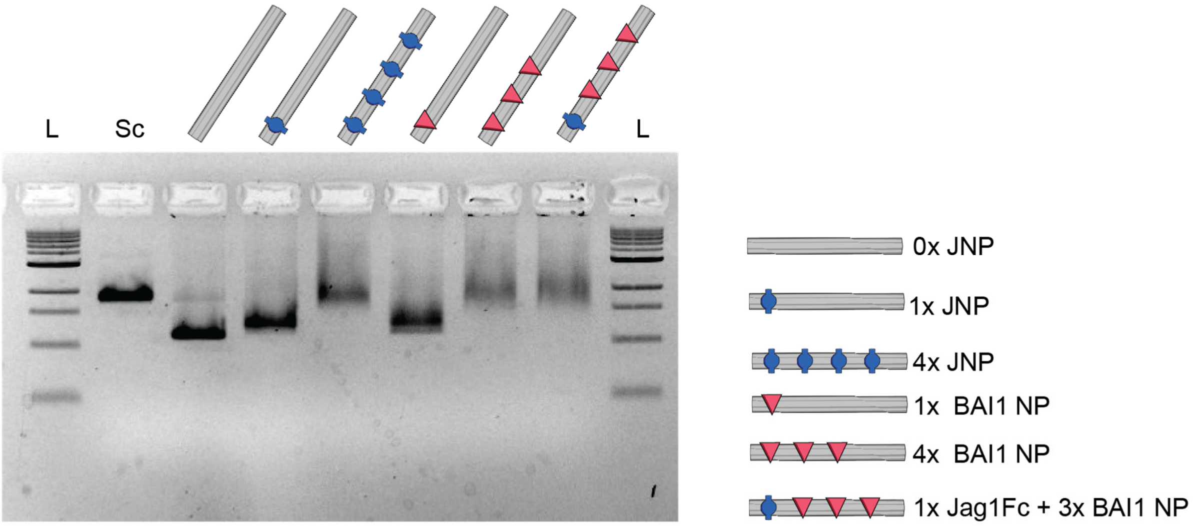


**Figure S5: Gel Retardation assays. A Coating of DNA nanostructures with oligolysine solution (K10).** Different ratios of azides (N) from oligolysine solution to phosphorus groups (P) of DNA origami were tested and run-on agarose gel stained with ethidium bromide. We observe a coating of DNA nanostructures occurred when more than 0,5:1 N:P ratio applied, since a mobility shift observed on the agarose gel. **B DNA nanopatterns decorated with Jag1 protein and/or BAI1 protein.** The two proteins have close molecular weights and similar charge under the same pH conditions. On the agarose gel stained with ethidium bromide we observe the same shift when one protein of Jag1 or BAI1 applied on NPs, as well as the same higher shift when four Jag1, four Bai1 or one Jag1+3BAI1 added on NPs.
